# Regulatory T cells and IFN-γ-producing Th1 cells play a critical role in the pathogenesis of Sjögren’s Syndrome

**DOI:** 10.1101/2024.01.23.576314

**Authors:** Yin-Hu Wang, Wenyi Li, Maxwell McDermott, Ga-Yeon Son, George Maiti, Fang Zhou, Anthony Tao, Dimitrius Raphael, Andre L. Moreira, Boheng Shen, Martin Vaeth, Bettina Nadorp, Shukti Chakravarti, Rodrigo S. Lacruz, Stefan Feske

## Abstract

**Objectives:** Sjögren’s Disease (SjD) is an autoimmune disorder characterized by progressive dysfunction, inflammation and destruction of salivary and lacrimal glands, and by extraglandular manifestations. Its etiology and pathophysiology remain incompletely understood, though a role for autoreactive B cells has been considered key. Here, we investigated the role of effector and regulatory T cells in the pathogenesis of SjD.

**Methods:** Histological analysis, RNA-sequencing and flow cytometry were conducted on glands, lungs, eyes and lymphoid tissues of mice with regulatory T cell-specific deletion of stromal interaction proteins (STIM) 1 and 2 (*Stim1/2^Foxp3^*), which play key roles in calcium signaling and T cell function. The pathogenicity of T cells from *Stim1/2^Foxp3^*mice was investigated through adoptively transfer into lymphopenic host mice. Additionally, single-cell transcriptomic analysis was performed on peripheral blood mononuclear cells (PBMCs) of patients with SjD and control subjects.

**Results:** *Stim1/2^Foxp3^* mice develop a severe SjD-like disorder including salivary gland (SG) and lacrimal gland (LG) inflammation and dysfunction, autoantibodies and extraglandular symptoms. SG inflammation in *Stim1/2^Foxp3^* mice is characterized by T and B cell infiltration, and transcriptionally by a Th1 immune response that correlates strongly with the dysregulation observed in patients with SjD. Adoptive transfer of effector T cells from *Stim1/2^Foxp3^* mice demonstrates that the SjD-like disease is driven by interferon (IFN)-γ producing autoreactive CD4^+^ T cells independently of B cells and autoantiboodies. scRNA-seq analysis identifies increased Th1 responses and attenuated memory Treg function in PBMCs of patients with SjD.

**Conclusions:** We report a more accurate mouse model of SjD while providing evidence for a critical role of Treg cells and IFN-γ producing Th1 cells in the pathogenesis of SjD, which may be effective targets for therapy.

## INTRODUCTION

Sjögren’s Disease (SjD) is a common autoimmune disorder with a prevalence of 0.5-1% in the general population. The clinical hallmarks of SjD are dry eyes and mouth (Sicca syndrome) due to lacrimal gland (LG) and salivary gland (SG) dysfunction, inflammation and destruction^1^. In addition to pathology of the exocrine glands, SjD is variably manifested by arthritis, vasculitis, neuropathy, pneumonitis, and it may be marked by dysfunction of the gastrointestinal tract, kidney and hematopoietic system. In some patients, the systemic disease becomes life-threatening, with the leading causes of death being interstitial lung disease (ILD), renal failure, cryoglobulinemic vasculitis and B cell lymphoma^2,3^. Patients with SjD have a 5-10% lifetime risk of non-Hodgkin lymphoma, which is 15-20 times higher than in the general population^4^. The pathophysiology of SjD remains incompletely understood, and clinical trials of immunomodulatory drugs that were successful in other autoimmune diseases showed mixed results in cases of SjD^4^.

SjD is associated with autoantibodies, including anti-nuclear antibodies (ANA), anti-Ro/SS-A and anti-La/SS-B antibodies^1,2^. Autoreactive B cells and autoantibodies are considered to be crucial for the pathogenesis of SjD^5,6^, which is supported by the promising outcomes of clinical trials focused on B cell depletion or inhibition of B cell activation^7,8^. Growing evidence suggests that T cells not only promote B cell activation and thus, contribute to the pathogenesis of SjD, but they also play an independent role in salivary and lacrimal glands inflammation and dysfunction in SjD. For instance, desiccating stress induces keratoconjunctivitis and lacrimal gland inflammation in animal models of SjD, and the disease can be transferred to non-stressed nude mice by injection of CD4^+^ T cells^9^. Lacrimal gland inflammation can also be induced in NOD-SCID mice by transferring CD8^+^ T cells from NOD mice^10^. Moreover, deletion of CD8^+^ T cells significantly alleviates gland inflammation and hyposalivation in two different murine SjD models; namely, NOD and *Il12p40^-/-^Il2ra^-/-^*mice^10,11^. In addition, mice lacking inhibitor of DNA binding 3 (ID3) develop many features typical of SjD, including sicca symptoms and gland inflammation, as well as anti-Ro and anti-La antibodies. Transfer of CD3^+^ T cells, but not B cells, from *Id3*-deficient mice induce sicca syndrome in sublethally irradiated host mice^12^. The M3 muscarinic acetylcholine receptor (M3R), which is highly expressed in the SG and induces saliva secretion^13^, is an autoantigen in SjD^14^. Lymphopenic *Rag1^-/-^* mice receiving CD3^+^ T cells from donor mice immunized with M3R peptides develop sialadenitis and hyposalivation^15^.

T regulatory (Treg) cells represent a distinct subset of CD4^+^ T cells that are immunosuppressive and maintain immunological self-tolerance and homeostasis^16^. The majority of Treg cells are characterized by the expression of the transcription factor Foxp3, but Foxp3-negative Treg cells, including Tr1 cells, also exist^17^. Foxp3^+^ Treg cells either develop during the negative selection of CD4^+^ T cells in the thymus or differentiate from conventional naive CD4^+^ T cells in peripheral tissues^18^. Thymus-derived Treg cells can further differentiate into specialized effector Treg subsets, including tissue-resident, memory-like Treg cells that acquire a tissue-specific gene expression program and maintain immune homeostasis in many organs^19,20^, as well as T follicular regular (Tfr) cells that shape the quality and quantity of humoral immune responses^21-23^. Defects in Treg development and/or function are associated with the pathogenesis of several autoimmune diseases, including type 1 diabetes (T1D), systemic lupus erythematosus (SLE), multiple sclerosis (MS) and rheumatoid arthritis (RA)^24^. The role of Treg cells in the pathogenesis of SjD remains ambiguous. The frequency of Treg cells in the blood of patients with SjD has alternatively been reported to be increased, decreased or unchanged compared to controls in various studies^25-34^. Data regarding Treg cells in SG tissues are also controversial. Whereas one study reported decreased Treg numbers in SG biopsies of patients with SjD compared to non-immune parotitis biopsies^25^, other groups showed a positive correlation between Treg infiltration and the severity of histological lesions^30,32^. In mouse models of SjD, Treg cells appear to prevent SjD-like disease. Mice with genetic deletion of IL-2 or the high-affinity IL-2 receptor α chain (IL-2Rα or CD25), which are critical for Treg function, spontaneously develop lymphocytic and neutrophilic SG and LG inflammation and dysfunction, whereas Scurfy mice, which carry a missense mutation in the *Foxp3* gene, only developed gland inflammation and loss of function upon LPS stimulation^35^. Although the study provided valuable insights into the involvement of Treg cells in SjD, it did not investigate the presence of autoantibodies, a key diagnostic feature of SjD^36^, or an in-depth immunological and molecular characterization of SG and LG inflammation.

Store-operated Ca^2+^ entry (SOCE) is an essential Ca^2+^ signaling pathway in T cells^37^. SOCE is initiated by T cell receptor (TCR) activation, which leads to the depletion of Ca^2+^ stores in the endoplasmic reticulum (ER). ER Ca^2+^ levels are sensed by stromal interaction molecules 1 and 2 (STIM1 and STIM2), and their reduction triggers the binding of STIM1 and STIM2 to ORAI1, which is located in the plasma membrane and forms the Ca^2+^ release-activated Ca^2+^ (CRAC) channel. The opening of ORAI1^+^ CRAC channels results in SOCE and induces Ca^2+^ signaling in T cells^38,39^. Patients with loss-of-function mutations in *STIM1* or *ORAI1* suffer from a combined immunodeficiency with infections and autoimmunity^38,40,41^. In mice, T cell-specific deletion of *Stim1* and *Stim2*, or *Orai1* and its homolog *Orai2*, abolishes SOCE and suppresses adaptive immune responses to infection. Moreover, the development and function of thymic-derived and peripheral Treg cells is compromised, resulting in multiorgan autoimmunity. This phenotype is exacerbated in mice with Treg-specific deletion of *Stim1* and *Stim2*, as male mice hemizygous for *Stim1^fl/fl^ Stim2^fl/fl^ Foxp3^YFPcre^*(abbreviated *Stim1/2^Foxp3^*) develop a scurfy-like disease. Autoimmunity in these mice is characterized by reduced frequencies of tissue-resident Treg cells and Tfr cells, as well as by the presence of a broad spectrum of autoantibodies^42^.

Here we show that ablation of SOCE in Treg cells in *Stim1/2^Foxp3^*mice results in a phenotype that recapitulates all the clinical features of SjD in human patients, including SG and LG inflammation with hyposalivation and reduced tear production, keratoconjunctivitis and anti-Ro and anti-La autoantibodies. Additional symptoms in mice that are often found in patients with SjD (although not part of the 2016 ACR/EULAR classification criteria for SjD^36^) include interstitial lung disease. SG inflammation in *Stim1/2^Foxp3^* mice is characterized by a Th1/Th2-mediated immune response and differential gene expression signatures like those found in SGs of patients with SjD^43,44^. The transfer of conventional CD4^+^ T cells from *Stim1/2^Foxp3^* mice was sufficient to induce severe SG and LG inflammation and dysfunction in lymphopenic host mice. The ability of potentially autoreactive CD4^+^ T cells to induce SjD-like disease was dependent on their ability to produce IFN-γ, whereas IFN-γ treatment of wild-type SG cells suppressed Ca^2+^ influx. These findings in *Stim1/2^Foxp3^* mice correlated with enhanced Th1 gene expression signatures and type I and II interferon responses in peripheral blood mononuclear cells (PBMCs) of patients with SjD. Moreover, we identified attenuated expression of genes in memory Treg cells of patients with SjD that are associated with Treg function. Together, our findings in mice and human patients provide evidence for a critical role of Treg cells and IFN-γ producing Th1 cells in the pathogenesis of SjD.

## RESULTS

### Loss of SOCE in Treg cells causes a SjD-like disease in mice

Targeted deletion of *Stim1* and *Stim2* genes in Treg cells of *Stim1/2^Foxp3^* mice abolishes SOCE (**Figure 1a**). Stimulation of CD4^+^CD25^hi^ Treg cells from spleen and lymph nodes (LNs) of hemizygous female *Stim1/2^Foxp3^* mice with the sacro-endoplasmic reticulum ATPase (SERCA) blocker thapsigargin (TG) resulted in similar depletion of ER Ca^2+^ stores compared to Foxp3-Cre-negative littermate controls, whereas Ca^2+^ levels following addition of extracellular Ca^2+^ to induce SOCE were almost completely abolished in *Stim1/2*-deficient Treg cells. We previously reported that male *Stim1/2^Foxp3^* mice, which lack STIM1/2 proteins in FoxP3^+^ Treg cells, and thus SOCE in these cells, develop multi-organ inflammation that is associated with the production of antibodies against a range of auto-antigens^42^. Because inflammation in *Stim1/2^Foxp3^*mice also included severe blepharitis with eyelid crusting^42^ and enlarged cervical lymph nodes (cLNs), we investigated if deletion of SOCE in FoxP3^+^ Treg cells results in inflammation of lacrimal and salivary glands. We observed massive lymphocytic infiltration of submandibular salivary glands in *Stim1/2^Foxp3^* mice but not in Cre-negative littermate control mice (Cre^-^ control), which was quantified using a modified focus score **(Figure 1b)**.

**Figure 1.**
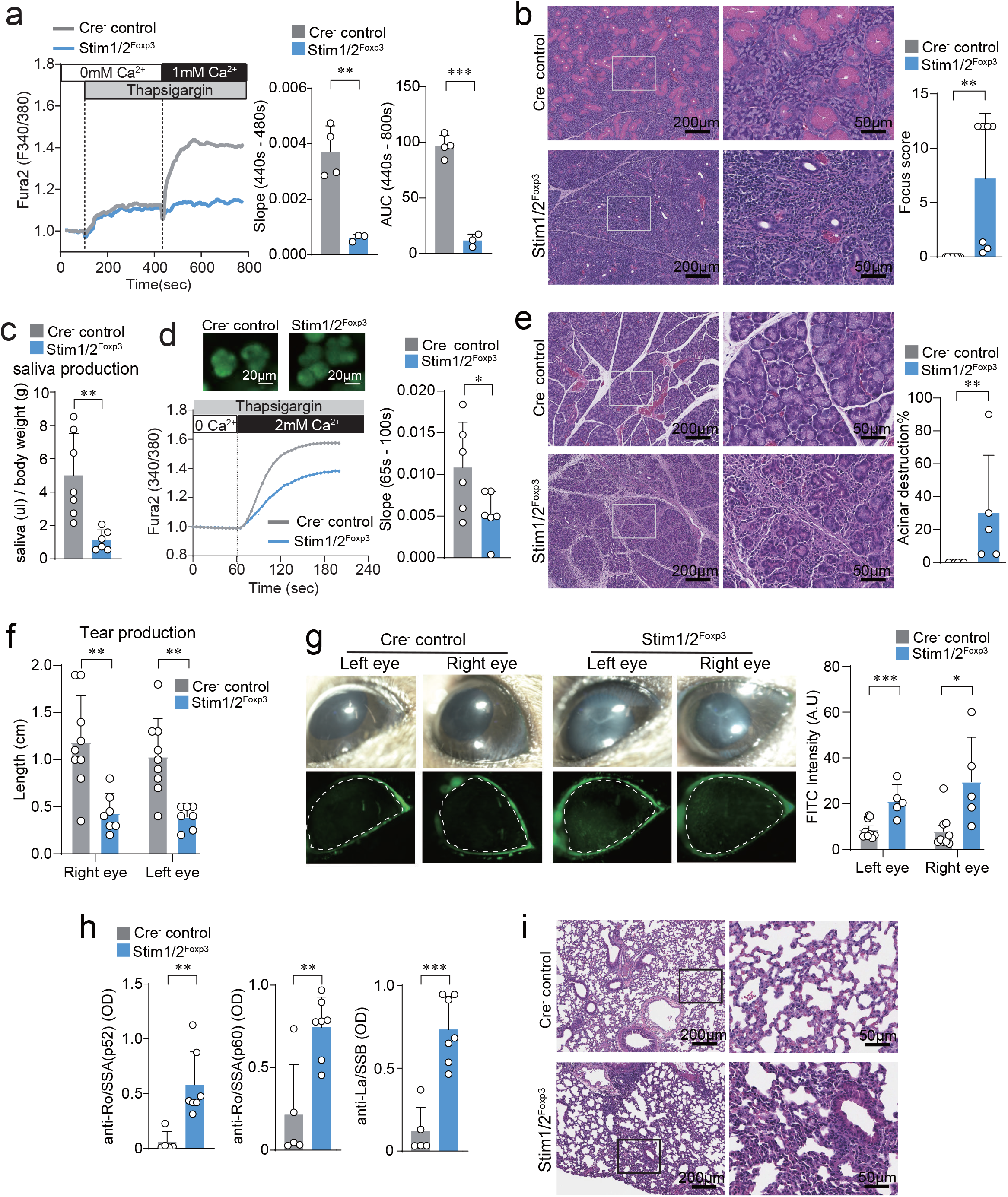
Loss of SOCE in Treg cells results in a SjD-like phenotype in mice. **a)** Analysis of store-operated Ca^2+^ entry (SOCE) in CD4^+^CD25^hi^ YFPcre-positive and YFPcre-negative control Treg cells isolated from spleen and lymph nodes (LNs) of female heterozygous *Stim1/2^Foxp3^* mice. Data are from 4 mice with 1 technical replicate per mouse. Fura-2 loaded cells were stimulated with thapsigargin (TG) in Ca^2+^-free Ringer solution followed by re-addition of 1 mM Ca^2+^. Slope of SOCE and area under the curve (AUC) at the indicated time points were measured. **b-i)** Analysis of exocrine gland inflammation, function and SjD-like phenotype in male *Stim1/2^Foxp3^*mice and Cre-negative littermate controls. **b)** Representative H&E staining and modified focus score of submandibular salivary glands (SMG). Data are from 7-8 mice per genotype with 1 gland (right) per mouse. **c)** Saliva production measured for 20 min after pilocarpine stimulation. Data are from 6-7 mice per cohort with 1 measurement per mouse. **d)** Analysis of SOCE in acini isolated from SMG of mice. Fura-2 loaded acini (top) were stimulated with TG in Ca^2+^-free Ringer solution followed by re-addition of 2 mM Ca^2+^. Bar graphs show slope of Ca^2+^ influx between 65s and 100s. Data are from 6 mice per cohort with 41-201 acini per mouse. **e)** Representative H&E staining and quantified destruction of exorbital lacrimal glands (ELG). Data are from 5-6 mice per cohort. **f)** Analysis of tear production by phenol red thread test. Data are from 7-9 mice per cohort. **g)** Representative ocular surface images and corneal fluorescein (FITC) staining. The bar graph shows the cumulative FITC intensity of right and left eye. Data are from 5-9 mice per cohort with 1 measurement per eye. **h)** Analysis of anti-SSA and anti-SSB autoantibodies concentrations in the serum of 5-7 mice per cohort by ELISA with 1 technical replicate per mouse. **i)** Representative H&E staining of lung tissues from 3 mice per cohort with 1 lung lobe (inferior) per mouse. Statistical analysis in all panels was conducted by two-tailed, unpaired Student’s *t*-test, and shown as means ± SD. \**P* < 0.05, \*\**P* < 0.01 and \*\*\**P* < 0.001.

To determine the effects of inflammation on SG function, we measured pilocarpine-induced saliva production. Saliva volumes were significantly lower in male *Stim1/2^Foxp3^* mice compared to Cre-negative littermate controls **(Figure 1c),** indicating impaired SG gland function. As Ca^2+^ signaling is critical for salivary gland function and SOCE was shown to be an important Ca^2+^ influx pathway in SG acinar cells^45,46^, we analyzed the effects of inflammation on SG function. Acini isolated from submandibular SG of male *Stim1/2^Foxp3^* mice and littermates were loaded with Fura-2 and stimulated with TG followed by addition of 2 mM extracellular Ca^2+^ **(Figure 1d)**. Acinar cell-inflamed SGs of *Stim1/2^Foxp3^* mice had significantly less SOCE compared to acini from uninflamed SGs of control mice. Of note, the structure of isolated acini from *Stim1/2^Foxp3^* mice was like those of control mice. Moreover, *Foxp3* is not expressed in SG cells and Foxp3-Cre does not delete *Stim1/2* genes in acinar cells, suggesting that the lower SOCE in SG acinar cells of the mutant mice is likely caused by a functional defect secondary to SG inflammation.

Given the macroscopic blepharitis observed in *Stim1/2^Foxp3^*mice, we next investigated if lacrimal glands are inflamed in these animals. Exocrine LGs from male *Stim1/2^Foxp3^* mice had profound inflammation, structural damage and severe fibrosis, which resulted in significantly lower gland areas with intact acini compared to the Cre-negative littermate controls **(Figure 1e)**. The analysis of lacrimal gland function using a phenol red thread test showed significantly lower tear production in male *Stim1/2^Foxp3^* mice compared to littermate controls **(Figure 1f)**. Reduced tear production and ocular dryness results in keratoconjunctivitis in human patients with SjD, which is one of the classification criteria for the disease^36^. We used fluorescein dye to detect epithelial abrasion at the ocular surface, because the dye it is taken up at sites of damage to the ocular surface glycocalyx and epithelial tight junctions^47^. The eyes of male *Stim1/2^Foxp3^* mice had significantly greater FITC fluorescence intensity compared to littermate controls **(Figure 1g),** suggesting a loss of epithelial cells and subsequent corneal damage due to LG dysfunction and ocular dryness^48^.

The breakdown of immunological tolerance in SjD, which is considered to be a crucial step in disease pathogenesis, is evident from the presence of pathognomonic auto-antibodies against Ro/SSA and La/SSB antigens^5,6,36^. To determine if SG and LG inflammation and dysfunction in *Stim1/2^Foxp3^* mice is associated with SjD-specific autoantibodies, we analyzed their presence in the serum of mice by ELISA. We detected significantly greater titers of anti-Ro/SSA (p52), anti-Ro/SSA(p60) and anti-La/SSB auto-antibodies in the serum of male *Stim1/2^Foxp3^* mice compared to Cre-negative littermate controls **(Figure 1h)**. Extraglandular manifestations associated with lymphocytic infiltration of the kidney or lung are common in many patients with SjD although they are not official ACR/EULAR classification criteria for the disease. Pulmonary involvement affects 16% of patients with primary SjD, and more than 45% of them were found to have interstitial pneumonia^49^. We observed extensive perivascular and peribronchiolar lymphocytic infiltration in the lungs of *Stim1/2^Foxp3^*mice but not in control mice **(Figure 1i)**. In addition, *Stim1/2^Foxp3^* mice showed focal to moderate interstitial pulmonary inflammation, with 1/3 of animals meeting the criteria for classification of lymphoid interstitial pneumonitis (LIP) with lymphocytic infiltrates in the alveolar walls associated with bronchitis and bronchiolitis with lymphoid aggregates^50,51^. Taken together, our findings demonstrate that *Stim1/2^Foxp3^*mice represent a new and more accurate animal model of SjD that satisfies all clinical diagnostic criteria of the human disease^36^ and recapitulates some of the typical extraglandular manifestations of SjD, including interstitial pneumonia.

### SOCE in Treg cells prevents lymphocytic salivary and lacrimal gland infiltration

A positive focus score (more than 50 infiltrating mononuclear immune cells per 4 mm^2^ SG section) is an important criterion for the diagnosis of SjD^36^. T and B lymphocytes represent the vast majority (more than 90%) of these cells in the minor SGs (MSGs) of patients with SjD but the composition of infiltrating immune cells changes with disease severity^52^. The frequencies of CD4^+^ T cells, Treg cells, B cells and macrophages (but not CD8^+^ T cells or NK cells) were significantly different in MSGs of patients with SjD with mild, intermediate or severe inflammatory lesions. We therefore investigated the composition of lymphocytes in the SGs, LGs and draining cervical LNs (cLNs) of male *Stim1/2^Foxp3^* mice and Cre-negative littermate controls. We found significantly higher frequencies and total numbers of CD138^high^B220^low^ plasma cells (PCs) and CD95^high^GL7^high^ germinal center B cells in the cLNs of *Stim1/2^Foxp3^* mice **(Figure 2a, b)**, suggesting enhanced B cell activation and expansion consistent with the greater autoantibody levels in these mice **(Figure 1h)**. There was also significantly higher frequencies and total numbers of both CD4^+^ and CD8^+^ T cells in the cLNs of *Stim1/2^Foxp3^* mice compared to Cre-negative littermates **(Figure 2c)**. Of note, the numbers of Foxp3^+^ Treg cells were also significantly higher in *Stim1/2^Foxp3^* mice compared to Cre-negative littermates **(Figure 2d)**. Because of their abolished SOCE, these Treg cells are dysfunctional, as we had reported previously^42^, explaining the severe SjD-like phenotype of *Stim1/2^Foxp3^* mice despite greater Treg numbers. In the submandibular SGs of *Stim1/2^Foxp3^* mice, we found significantly greater frequencies and numbers of CD4^+^ and CD8^+^ T cells, as well as CD19^+^ B cells compared to Cre-negative littermates **(Figure 2e, f)**. A similarly massive difference was observed in LGs of *Stim1/2^Foxp3^* mice compared to Cre-negative littermates **(Figure 2g)**. A sizable percentage of CD4^+^ T cells in the SGs of *Stim1/2^Foxp3^* mice produced IFN-γ and TNF-α, which was even more pronounced in LG-infiltrating CD4^+^ T cells **(Supplementary Figure 1a, b),** suggesting that CD4^+^ T cells in SGs and LGs are activated. These findings are consistent with the increased IFN-γ and TNF-α levels found in the serum, saliva and SGs of patients with SjD^53-58^. Of note, we were not able to detect significant numbers of Treg cells in the SGs of either *Stim1/2^Foxp3^* or littermate control mice (data not shown). Together, our data demonstrate that SG and LG inflammation in *Stim1/2^Foxp3^*mice is dominated by CD4^+^ and CD8^+^ T cells, which is similar to that observed in glands of patients with SjD^52^.

**Figure 2.**
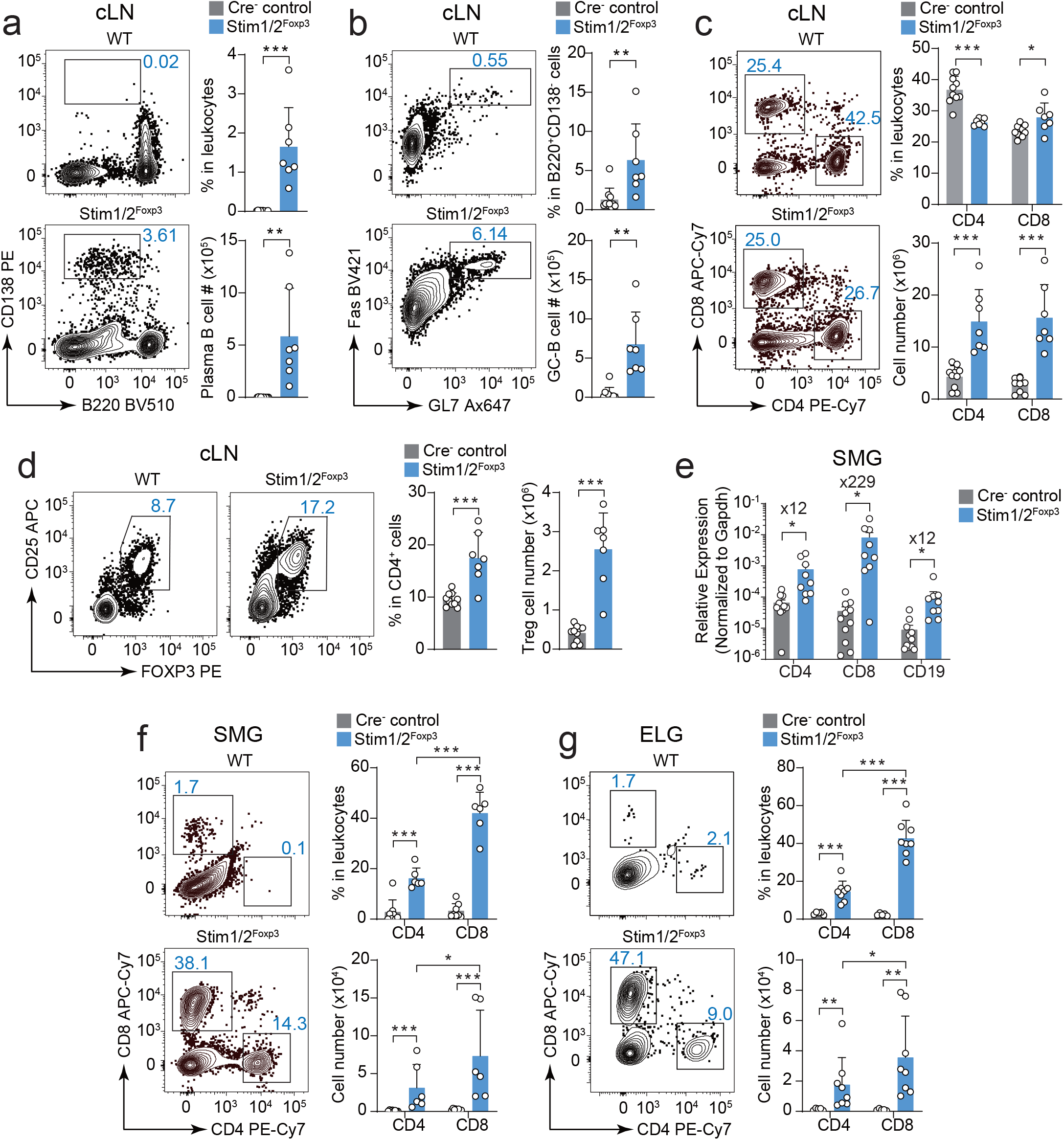
SOCE in Treg cells prevents lymphocytic infiltration of salivary and lacrimal glands. Analysis of immune cell populations in cervical lymph nodes (cLN) and exocrine glands of male *Stim1/2^Foxp3^* mice and Cre-negative littermate controls. **a-d)** Frequencies and absolute numbers of plasma cells **(a)**, germinal center B cells **(b)**, CD4^+^ and CD8^+^ T cells **(c)** and Foxp3^+^CD25^hi^ Treg cells **(d)** in cLNs. Representative flow cytometry plots and bar graphs showing data from 7-10 mice per cohort with 1 technical replicate per mouse. **e)** Expression levels of *Cd4, Cd8, Cd19* mRNA in total submandibular gland (SMG) tissue measured by RT-qPCR and normalized to *Gapdh* mRNA levels. Data are from 9-11 mice per cohort with 1 technical replicate per mouse. **(f)** Frequencies and absolute numbers of CD4^+^ and CD8^+^ T cells in SMG **(f)** and exorbital lacrimal gland (ELG, **g**). Representative flow cytometry plots and bar graphs from 5-8 mice per cohort with 1 technical replicate per mouse. All data were analyzed by two-tailed, unpaired Student’s *t* test, and shown as means ± SD. \**P* < 0.05, \*\**P* < 0.01 and \*\*\**P* < 0.001.

### Loss of SOCE in Treg cells promotes Th1- and Th2-mediated inflammation of salivary glands

To gain insights into the causes of SG inflammation and damage in *Stim1/2^Foxp3^* mice, we conducted RNA-sequencing (RNA-seq) of submandibular SGs of male *Stim1/2^Foxp3^* mice and *Foxp3*-YFPcre^-^ littermates. The transcriptional analysis identified 2044 up- and 1152 down-regulated genes in the SGs of *Stim1/2^Foxp3^* mice **(Figure 3a)**. An unbiased pathway enrichment and network analyses of differentially expressed genes (DEGs) revealed that most enriched pathways in the SGs of *Stim1/2^Foxp3^* mice were related to T cell activation, TCR signaling, chemokines & immune cell migration and other immune-related processes **(Figure 3b)**. Of note in this context the expression of markers for acinar and ductal SG structures, such as the water channel aquaporin 5 (*Aqp5*) or different keratins^59^, was not significantly diminished **(Supplementary Figure 2a-c)**. We further confirmed normal AQP5 levels and distribution in the SG of *Stim1/2^Foxp3^*mice by immunofluorescence **(Supplementary Figure 2d)**, suggesting that the SjD-like phenotype of male *Stim1/2^Foxp3^* mice is not due to altered expression of SG genes, but rather SG dysfunction caused by SG inflammation. Further functional pathway enrichment analyses of RNA-seq data using IPA (Ingenuity Pathway Analysis) and KEGG (Kyoto Encyclopedia of Genes and Genomes) databases identified T helper (Th) 1 and Th2-related pathways to be among the most significantly dysregulated pathways **(Figure 3c)**. Gene Set Enrichment Analysis (GSEA) of DEGs in SGs of *Stim1/2^Foxp3^*mice moreover showed a significant enrichment of genes related to type I IFN production and downstream IFNα/β signaling **(Figure 3d)**, which is reminiscent of the increased type I IFN levels and upregulated expression of type I IFN-stimulated genes found in exocrine glands and the blood of patients with SjD^60,61^.

**Figure 3.**
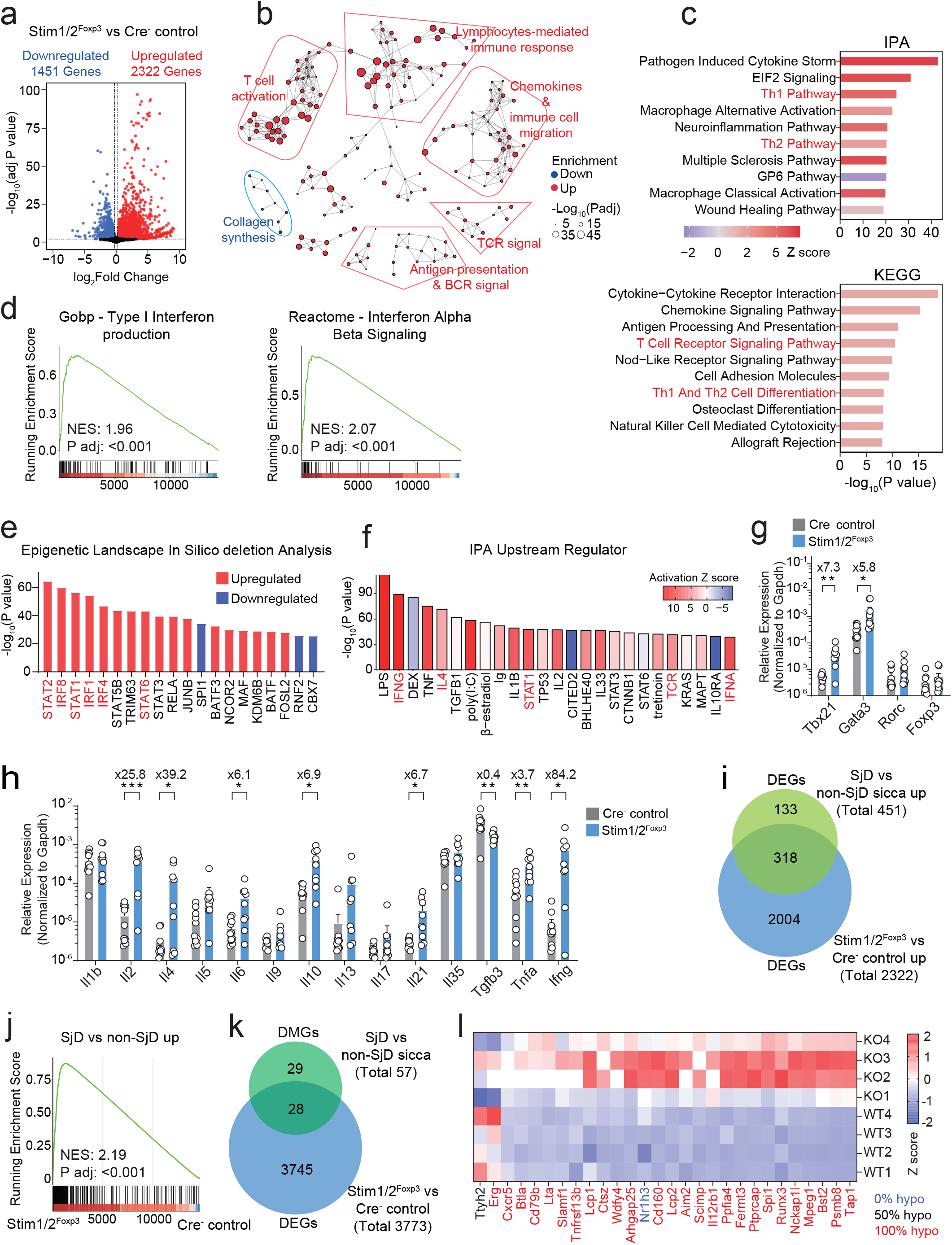
Loss of SOCE in Treg cells results in Th1/Th2-dominant inflammation of salivary glands. **a–f)** RNA-sequencing and transcriptome analysis of SMG isolated from male *Stim1/2^Foxp3^* mice and Cre-negative littermate controls (4 mice per cohort with 1 gland (right) per mouse). **a)** Differentially expressed genes (DEG). **b)** Pathway network analysis based on DEGs. Downregulated and upregulated pathways in SMG of *Stim1/2^Foxp3^*mice are marked in blue and red, respectively. **c)** Pathway enrichment analysis based on DEGs in SMG. Top 10 dysregulated pathways identified by analysis of IPA and KEGG databases are ranked by *P* value (-log_10_). **d)** Gene set enrichment analysis (GSEA) of GOBP Type I Interferon and Reactome Interferon alpha beta signaling pathways in SMG of *Stim1/2^Foxp3^* vs control mice. NES, normalized enrichment score. **e, f)** Epigenetic landscape in silico deletion analysis (LISA, in **e**) and IPA upstream regulator analysis **(f)** of DEGs ranked by *P* value. Down- and upregulated upstream factors in SMG of *Stim1/2^Foxp3^* mice are indicated in blue and red, respectively. **g, h)** mRNA expression levels of lineage specific transcription factors of Th1, Th2, Th17 and Treg cells **(g)** and cytokines **(h)** in SMG measured by RT-qPCR and normalized to *Gapdh* mRNA levels. Data are from 9 (*Stim1/2^Foxp3^*) and 11 (control) mice with 1 technical replicate per mouse. **i, j)** Analysis of dysregulated gene expression in salivary glands of patients with SjD and *Stim1/2^Foxp3^* mice. **i)** Comparison of upregulated genes in SMG of male *Stim1/2^Foxp3^* vs. Cre-negative littermate control mice and parotid and labial glands of patients with SjD vs. non-SjD Sicca^44^. **j)** GSEA of upregulated DEG in *Stim1/2^Foxp3^*mice against upregulated DEG in patients with SjD^44^. **k, l)** Comparison of upregulated genes in SMG of male *Stim1/2^Foxp3^*vs. Cre-negative littermate control mice to differentially methylated genes (DMG) in the labial SG of patients with SjD^43^. Red and blue color indicates high and low expression calculated by Z score per column. The degree of hypomethylation for each gene is indicated by the font color (blue, 0%; black, 50%; red, 100%). Statistical analysis in **g** and **h** by two-tailed, unpaired Student’s *t*-test. Data in all other panels were analyzed using Wilcoxon rank-sum test, and significance was adjusted using the Benjamini-Hochberg method. Results are shown as means ± SD. \**P* < 0.05, \*\**P* < 0.01 and \*\*\**P* < 0.001.

To identify upstream regulators of altered gene expression programs in SGs of *Stim1/2^Foxp3^* mice, we conducted IPA upstream regulator and Epigenetic Landscape In Silico deletion Analysis (LISA)^62^. LISA, which queries transcriptional regulators based on chromatin immunoprecipitation sequencing (ChIP-Seq) datasets^62^, identified *Stat1, Stat2, Irf1* and *Irf8*, as well as *Stat6, Batf* and *Irf4*, which regulate Th1 and Th2 differentiation and function, respectively^63^ **(Figure 3e)**. An IPA upstream regulator analysis, which in addition to transcription factors includes signaling molecules, cytokines and extrinsic modulators of cell function, identified many Th1- and Th2-related cytokines and transcription factors, such as IFN-γ, TNF, IL-4 and STAT1, as regulators of DEGs in SGs of *Stim1/2^Foxp3^* mice **(Figure 3f)**. To confirm the Th1 and Th2 signatures identified by RNA-seq, we analyzed the mRNA expression of Th1, Th2, Th17 and Treg-defining transcription factors (TFs) and cytokines in submandibular SGs by RT-qPCR. The expression of *Tbx21* (Th1) and *Gata3* (Th2) was significantly greater in SGs of male *Stim1/2^Foxp3^* mice compared to Cre-negative littermate controls, whereas *Rorc* (Th17) and *Foxp3* (Treg) levels were comparable **(Figure 3g)**. Moreover, mRNA levels of the Th1 and Th2 cytokines, *Ifng* and *Il4*, respectively, were significantly greater *Stim1/2^Foxp3^* mice compared to Cre-negative controls, as were those of *Il2, Il10, Il21* and *Tnf* **(Figure 3h)**.

The greater Th1 and Th2 signatures in SGs of *Stim1/2^Foxp3^*mice compared to their Cre-negative controls are consistent with altered Th1 and Th2 cytokine levels reported in patients with SjD^44,54-58,64-66^. We compared the gene expression in SGs of *Stim1/2^Foxp3^*mice to those of patients with SjD by analyzing RNA-seq data from SGs of patients with SjD and patients with sicca but without SjD^44^. 451 DEGs were selectively upregulated in parotid and labial glands of patients with SjD, of which 318 were also upregulated in the SGs of *Stim1/2^Foxp3^* mice **(Figure 3i)** and GSEA of DEGs identified in SGs of human patients showed a significant enrichment of these genes in SGs of *Stim1/2*-deficient mice **(Figure 3j)**. A genome-wide DNA methylation study of labial SG biopsies of patients with SjD had identified an epigenetic signature that consisted of 57 differentially methylated genes (DMGs) that are associated with disease, suggesting that these genes are involved in the pathogenesis of SjD^43^. We found that 28 of these 57 DMGs are also differentially expressed in the SGs of *Stim1/2^Foxp3^* mice **(Figure 3k)**. The majority of these 28 genes were hypomethylated in patients (indicating their transcriptional activation) and expressed more highly in *Stim1/2^Foxp3^* mice compared to Cre-negative littermate controls **(Figure 3l)**. Collectively, these data show that changes in gene expression in SGs of mice due to impaired SOCE in FoxP3^+^ Treg cells correlate well with transcriptional and epigenetic changes in SGs of patients with SjD, thus pointing to a shared disease pathophysiology.

### Conventional CD4^+^ T cells from *Stim1/2^Foxp3^* mice are sufficient to induce SjD-like disease

Given the greater T cell numbers and Th1 and Th2 signatures in the SGs of the *Stim1/2^Foxp3^* mice compared to their Cre-negative littermates, we hypothesized that SG and LG inflammation is caused by autoreactive CD4^+^ T cells responding to a gland-specific autoantigen. To test this hypothesis, we isolated CD4^+^ CD25^-^ and CD8^+^ T cells from cLNs of male *Stim1/2^Foxp3^*mice and Cre-negative littermates and adoptively transferred them to lymphopenic *Rag1*^-/-^ mice **(Figure 4a, Supplementary Figure 3a)**. Host mice showed robust immune cell infiltration and greater frequencies of CD4^+^ T cells in their SGs 2-4 weeks after adoptive transfer of CD4^+^ T cells from *Stim1/2^Foxp3^* compared to control mice **(Figure 4b, c)**. All host mice injected with CD4^+^ T cells from *Stim1/2^Foxp3^* mice had significantly less saliva production compared to those injected with control CD4^+^ T cells **(Figure 4d)**. In addition to SGs, we found severe blepharitis of host mice that had received CD4^+^ T cells from *Stim1/2^Foxp3^* mice **(Figure 4e)**. Macroscopic ocular inflammation correlated well with immune cell infiltration and fibrosis of a substantial portion of LGs **(Figure 4f)**. Moreover, the numbers of CD4^+^ T cells were significantly greater in LGs of host mice injected with CD4^+^ T cells of *Stim1/2^Foxp3^* mice compared to host mice receiving control donor cells **(Figure 4g)**. Consequently, tear production was significantly impaired in the test mice compared to the control mice **(Figure 4h)**. Moreover, we observed severe conjunctival inflammation following transfer of CD4^+^ T cells from *Stim1/2^Foxp3^* mice **(Figure 4i)**. In parallel experiments, we injected *Rag1^-/-^* hosts with CD8^+^ T cells from *Stim1/2^Foxp3^* mice or Cre-negative controls because we had observed increased numbers of CD8^+^ T cells in the cLNs and SGs of *Stim1/2^Foxp3^* mice (**Figure 2**). The transfer of CD8^+^ T cells, however, did not cause SG or LG inflammation, had no effects on saliva or tear production and serum cytokine levels in host mice **(Supplementary Figure 3b-h)**. These data demonstrate that conventional CD4^+^ T cells isolated from cLNs of *Stim1/2^Foxp3^* mice are sufficient to cause SjD-like disease in the absence of B cells or autoantibodies.

**Figure 4.**
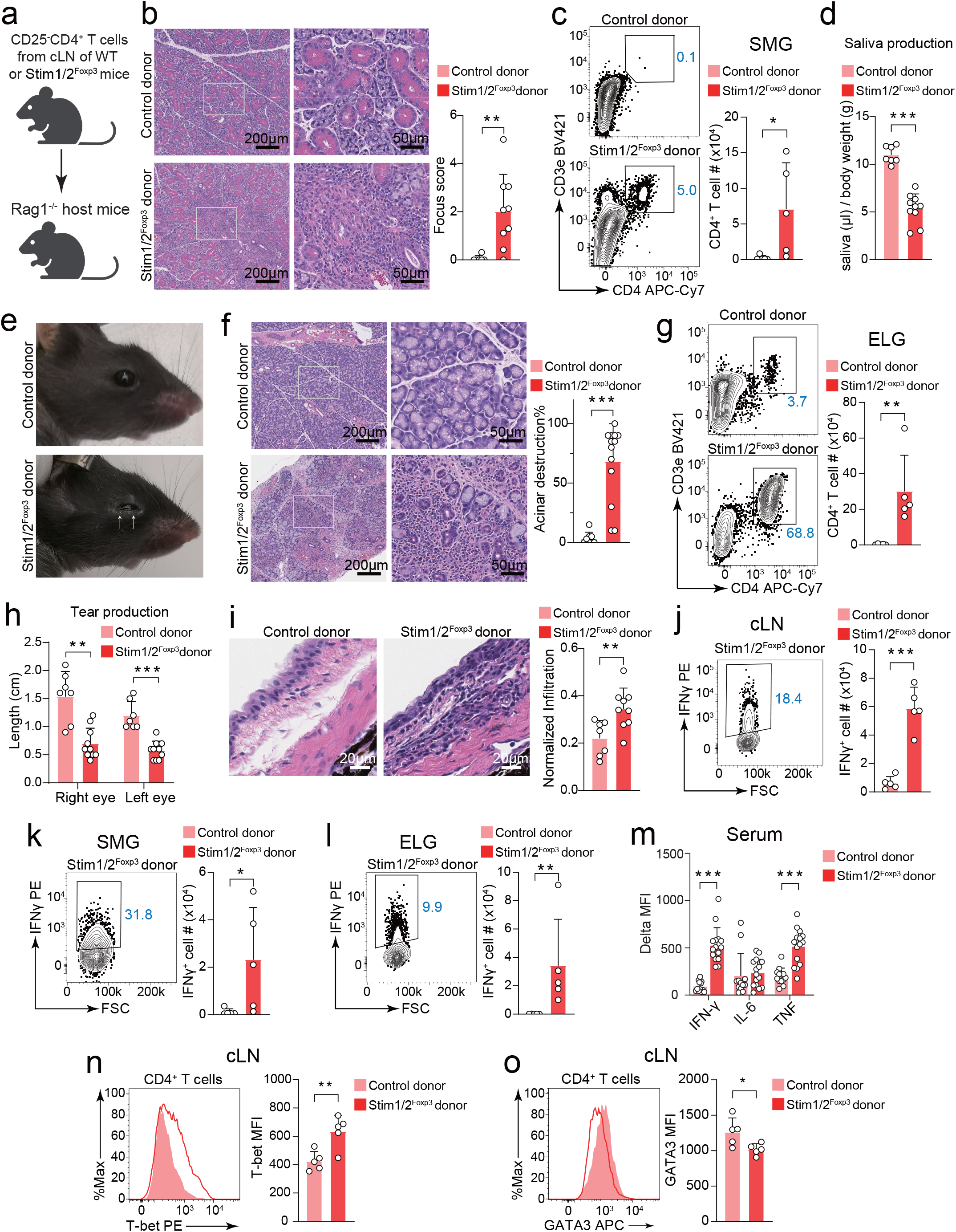
Autoreactive CD4^+^ T cells from *Stim1/2^Foxp3^* mice are sufficient to transfer SjD-like disease. **a)** Design of experiments shown in panels **b–o**. CD4^+^CD25^-^ T cells were isolated from cLN of male *Stim1/2^Foxp3^* and Cre-negative littermate control mice, and injected i.v. into male *Rag1^-/-^*mice. **b)** Representative H&E staining and modified focus scores of SMG from *Rag1^-/-^* host mice. Data are from 6-9 mice per cohort with 1 gland (right) per mouse. **c)** Representative flow cytometry plots numbers of CD4^+^ T cells in SMG of host mice. Data are from 5 mice per cohort with 1 technical replicate per mouse. **d)** Saliva production over a period of 20 min after pilocarpine stimulation of *Rag1^-/-^* host mice. Data are from 7-10 mice per cohort with 1 measurement per mouse. **e, f)** Representative ocular images, H&E staining and quantified exorbital lacrimal gland (ELG) destruction in *Rag1^-/-^* host mice. Data are from 8-13 mice per cohort with 1 eye or gland (right) per mouse. **g)** Representative flow cytometry plots and numbers of CD4^+^ T cells in ELG of host mice. Data are from 5 mice per cohort with 1 technical replicate per mouse. **h)** Analysis of tear production by phenol red thread test. Data are from 7-10 mice per cohort with 1 measurement per eye. **i)** Representative H&E staining of conjunctiva and quantified conjunctivitis from *Rag1^-/-^*host mice. Data are from 7-9 mice per cohort with 2 eyes per mouse. **j-l)** Representative flow cytometry plots and numbers of CD4^+^IFNγ^+^ T cells in cLN **(j)**, SMG **(k)** and ELG **(l)** of host mice. T cells were restimulated for 4h with PMA + Ionomycin. Data are from 5 mice per cohort with 1 technical replicate per mouse. **m)** Analysis of IFN-γ, TNF and IL-6 concentration in the serum of *Rag1^-/-^* host mice by CBA, Data are from 12 to 15 mice per cohort with 1 technical replicate per mouse. **n, o)** Representative histogram plots and means +/- SD of mean fluorescence intensities (MFI) of T-bet **(n)** and GATA3 **(o)** protein expression in donor CD4^+^ T cells isolated from cLN of host mice. Data are from 5 mice per cohort with 1 technical replicate per mouse. Statistical analyses were conducted by two-tailed, unpaired Student’s *t*-test, and shown as means ± SD. \**P* < 0.05, \*\**P* < 0.01 and \*\*\**P* < 0.001.

To understand how transferred CD4^+^ T cells cause disease, we analyzed their differentiation state and cytokine production. The transfer of conventional CD4^+^ T cells from *Stim1/2^Foxp3^*mice resulted in significantly greater IFN-γ production by donor CD4^+^ T cells in cLNs, SGs and LGs compared CD4^+^ T cells originating from littermate controls **(Figure 4j-l)**. Greater levels of IFN-γ were also detected in the serum of host mice receiving CD4^+^ T cells from *Stim1/2^Foxp3^* mice compared those injected with control CD4^+^ T cells **(Figure 4m)**. We also found greater serum concentrations of TNF, whereas levels of IL-6 were comparable. In contrast, we did not detect IL-4-producing T cells in cLNs and SGs, or IL-4 in the serum of host mice that had been injected with CD4^+^ T cells from *Stim1/2^Foxp3^*or control mice (data not shown). In addition to greater production of IFN-γ, CD4^+^ T cells from *Stim1/2^Foxp3^* mice isolated from cLNs of host mice 2-4 weeks after transfer expressed significantly higher levels of T-bet compared to control CD4^+^ T cells, whereas the expression of GATA3 was moderately decreased **(Figure 4n, o)**. These data demonstrate that the conventional CD4^+^ T cells from *Stim1/2^Foxp3^*mice which cause SjD-like disease in lymphopenic host mice are Th1 cells.

It is likely that conventional CD4^+^ T cells from *Stim1/2^Foxp3^*mice recognize SG- and LG-specific autoantigens that induce their activation and differentiation into Th1 cells. We analyzed the frequencies of TCR V beta chains (Vβ) in adoptively transferred CD4^+^ and CD8^+^ T cells isolated from the cLNs of *Rag1*^-/-^ mice to analyze their clonal expansion. We observed significantly higher frequencies of TCR Vβ13^+^CD4^+^ and TCR Vβ13^+^CD8^+^ donor T cells derived from *Stim1/2^Foxp3^* compared to littermate control mice **(Supplementary Figure 4a,b)**. To investigate if the ability of T cells to induce a SjD-like disease is dependent on Vβ13 expression, we isolated CD4^+^CD25^-^Vβ13^-^ and CD8^+^Vβ13^-^ T cells from cLNs of *Stim1/2^Foxp3^* mice and injected them into *Rag1*^-/-^ mice **(Supplementary Figure 4c)**. (Note that the frequencies of Vβ13^+^CD4^+^ and CD8^+^ T cells were too low to be used for adoptive transfer experiments). Recipient mice were analyzed four weeks after T cell transfer for SG and LG inflammation and function. Despite the absence of Vβ13^+^CD4^+^ and Vβ13^+^CD8^+^ T cells in cLNs of host mice **(Supplementary Figure 4d)**, we observed moderate lymphocytic infiltration of the SGs of host mice following transfer of Vβ13^-^ CD4^+^ T cells from *Stim1/2^Foxp3^* mice, whereas Vβ13^-^CD8^+^ T cells did not cause inflammation **(Supplementary Figure 4e)**. Vβ13^-^CD4^+^ T cells from *Stim1/2^Foxp3^* mice induced a moderate reduction in saliva production compared to Vβ13^-^CD8^+^ T cells from *Stim1/2^Foxp3^* mice **(Supplementary Figure 4f)** and CD4^+^ T cells from Cre-negative littermate controls **(Figure 4d)**. Vβ13^-^CD4^+^ T cells of *Stim1/2^Foxp3^*mice induced severe blepharitis, lymphocytic infiltration of LGs and significantly lower tear production compared to transfer of Vβ13^-^CD8^+^ T cells **(Supplementary Figure 4g-i)**. Taken together, these data indicate that although the frequencies of Vβ13^+^CD4^+^ T cells are higher in *Stim1/2^Foxp3^*mice, they are not required to cause SjD-like disease as injection of Vβ13^-^CD4^+^ T cells can cause a SjD-like disease in *Rag1*^-/-^ mice.

### IFN-γ expression is required for the ability of CD4^+^ T cells to cause SjD-like disease

Because of the dominant Th1 signature in cLNs and SGs of *Stim1/2^Foxp3^*mice, we hypothesized that Th1 cells and their ability to produce IFN-γ are essential to induce SjD-like disease in mice. To test this hypothesis, we deleted *Ifng* expression in conventional CD4^+^ T cells from *Stim1/2^Foxp3^* mice by transduction with shRNAs targeting *Ifng* (or *Cd19* as a control) *in vitro.* Transduced T cells were injected into *Rag1^-/-^* host mice and their effects analyzed 10-12 weeks later **(Figure 5a)**. Compared to sh*Cd19* controls, sh*Ifng*-transduced CD4^+^ T cells isolated from cLNs of host mice showed significantly lower IFN-γ production, whereas expression of TNF was normal **(Figure 5b)**. Deletion of *Ifng* did not affect the expression of T-bet or the chemokine receptor CXCR3, which is preferentially expressed on Th1 cells^67^, suggesting an intact Th1 differentiation **(Figure 5c, d)**. Likewise, expression of the Th2-specific transcription factor GATA3 was also normal **(Figure 5e)**. The analysis of submandibular and parotid SGs of host mice that had received either sh*Cd19* or sh*Ifng-*transduced CD4^+^ T cells showed a similar extent of immune cell infiltration **(Figure 5f)**. The production of saliva, however, was significantly impaired only in mice injected with sh*Cd19*-transduced CD4^+^ T cells, but normal in mice receiving *Ifng*-deficient CD4^+^ T cells **(Figure 5g)**. Similar observations were made regarding the inflammation and function of LGs. Whereas sh*Cd19*-transduced CD4^+^ T cells from *Stim1/2^Foxp3^* mice caused severe blepharitis, lymphocytic LG infiltration and a significant reduction of tear production, mice receiving *Ifng*-deficient CD4^+^ T cells showed no signs of LG dysfunction **(Figure 5h-j)**. These data demonstrate that the ability of autoreactive effector Th1 cells from *Stim1/2^Foxp3^* mice to cause SjD-like disease depends on their production of IFN-γ.

**Figure 5.**
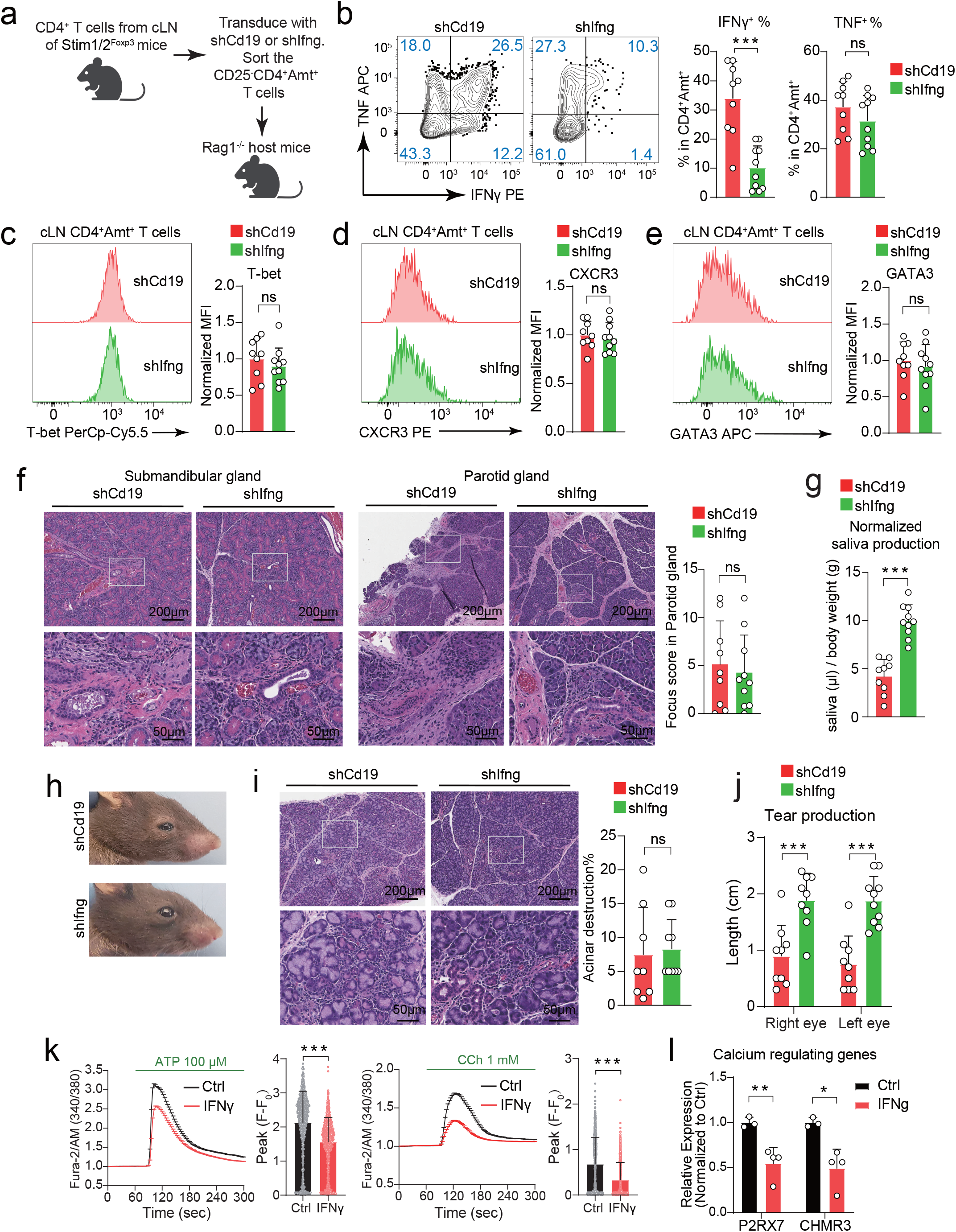
*Ifng* deletion in transferred autoreactive CD4^+^ T cells attenuates a SjD-like disease. **a)** Design of experiments shown in panels **b–j**. CD4^+^CD25^-^ T cells were isolated from cLN of male *Stim1/2^Foxp3^* and transduced with sh*Cd19* or sh*Ifng.* Transduced (Amt^+^) T cells were injected i.v. into male *Rag1^-/-^* mice. **b)** Representative flow cytometry plots and frequencies of IFNγ^+^CD4^+^Amt^+^ and TNF^+^CD4^+^Amt^+^ T cells in cLN of host mice that had received sh*Cd19* or sh*Ifng* transduced cells. T cells were stimulated with PMA + Ionomycin. **c–e)** Representative histogram plots and means +/- SD of mean fluorescence intensities (MFI) of T-bet **(c)**, CXCR3 **(d)**, and GATA3 **(e)** protein expression in donor T cells isolated from cLN of host mice. Data are from 2 independent experiment and 9-10 mice per cohort; MFI was normalized to the average value of the sh*Cd19* group in each experiment. **f)** Representative H&E staining and modified focus scores of SMG and Parotid gland (PG) from *Rag1^-/-^* host mice. **g)** Saliva production over a period of 20 min after pilocarpine stimulation of *Rag1^-/-^* host mice. **h, i)** Representative ocular images, H&E staining and quantified acinar destruction of ELG from *Rag1^-/-^*host mice. **j)** Analysis of tear production by phenol red thread test. Data in **(b–j)** are from 8-10 mice per cohort with 1 technical replicate or measurement, or 1 gland (right) per mouse. **k)** Ca^2+^ influx in human acinar cells (NS-SV-TT-AC) after 20 ng/ml IFNγ treatment for 4 days. Fura-2 loaded cells were stimulated with 100 μM ATP or 1 mM carbachol in 2 mM Ca^2+^ Ringer solution. Bar graphs show baseline (F_0_) normalized peak (F) Ca^2+^ levels. Data are from 3-4 independent experiments and 991–1,103 cells. **l)** *P2RX7* and *CHRM3* mRNA levels in human acinar cells treated with or without 20 ng/ml IFNγ for 4 days. RT-qPCR data were normalized to *Gapdh* housing control expression. Data are from 3-4 independent experiments per cohort with 3 technical replicates in each experiment. All data were analyzed by two-tailed, unpaired Student’s *t*-test, and shown as means ± SD. \**P* < 0.05, \*\**P* < 0.01 and \*\*\**P* < 0.001.

The deletion of *Ifng* in adoptively transferred CD4^+^ T cells prevented hyposalivation but had no effect on the extent of immune cell infiltration into SGs **(Figure 5f, g)**. These findings suggested that SG dysfunction may be due to the proinflammatory function of infiltrating Th1 cells; more specifically the effects of IFN-γ. The notion of Th1 cells and IFN-γ causing SG dysfunction is consistent with our earlier finding that SOCE was reduced in SG acinar cells from *Stim1/2^Foxp3^* **(Figure 1d)** and reports that IFN-γ decreases the proliferation and Ca^2+^ influx of human SG cells^68^. To test if IFN-γ affects SG function, we cultured human SG acinar cells in the absence or presence of IFN-γ for 4 days *in vitro* and measured Ca^2+^ signals as a read-out for their function. Treatment of SG acinar cells with IFN-γ resulted in significantly less Ca^2+^ influx induced by 100 μM ATP or 1 mM carbachol compared to control-treated cells **(Figure 5k)**.

Whereas ATP activates purinergic P2X7 receptors (encoded by *P2XR7*) and induces saliva secretion by the mouse submandibular glands^69^, carbachol activates the M3 subtype of muscarinic acetylcholine receptors (encoded by *CHRM3*), which is the main receptor involved in salivary flow^70^. To understand why ATP and carbachol-induced Ca^2+^ influx is reduced in IFN-γ treated SG cells, we analyzed the mRNA levels of *P2XR7* and *CHRM3.* IFN-γ treatment of human SG acinar cells resulted in significantly lower *P2XR7* and *CHRM3* expression levels compared to control-treated cells **(Figure 5l**). Collectively, these data provide a molecular mechanism by which IFN-γ suppresses Ca^2+^ influx and the function of SG acinar cells, and an explanation why deletion of *Ifng* in CD4^+^ T cells from *Stim1/2^Foxp3^* mice is protective against SG dysfunction.

### Greater IFN signaling and activity of Th1 cells occurs in patients with SjD

Given that *Stim1/2^Foxp3^* mice recapitulate most if not all manifestations of human SjD, we hypothesized that the underlying mechanisms of disease are similar in mice and patients. We therefore performed single-cell RNA-seq (scRNA-seq) of peripheral blood mononuclear cells (PBMCs) isolated from 9 patients with SjD and 8 patients with sialadenitis and sicca symptoms but without SjD **(Supplementary Table 3)**. All the patients with SjD except for one were anti-SSA/Ro^+^ and had ACR/EULAR scores ranging from 5-9, whereas controls were auto-antibody negative and had scores of 0 or 1. Analysis of the scRNA-seq data using a Louvain algorithm identified 7 clusters of cell types, including B cells, monocytes, NK cells, dendritic cells, CD4^+^ T cells and CD8^+^ T cells (**Supplementary Figure 5a**). Because CD4^+^ T cells had emerged as the main drivers of SjD-like disease in *Stim1/2^Foxp3^* mice, we focused our further analysis of the scRNA-seq data on CD4^+^ T cells. We identified a total of 12 different subsets of CD4^+^ T cells, including naïve, memory, Th1, Th2, Th17, CTL, Tfh, Tr1 and Treg cells, of which the naïve, memory and Treg populations could be further divided into 2 distinct populations **(Figure 6a)**. Neither cluster or cell type identified was unique to the patients with SjD or the controls **(Supplementary Figure 5b)**. To determine the composition of the CD4^+^ compartment in the patients with SjD, we normalized the frequency of T cell subsets to the total number of CD4^+^ T cells per patient and calculated their mean occurrence. The frequencies of all identified CD4^+^ T cell subsets was comparable in the patients with SjD and the controls except for *FOXP3^-^ LAG3^+^IL10^high^CTLA4^+^* type 1 regulatory (Tr1) cells, which were more abundant in the patients with SjD **(Figure 6b, Supplementary Figure 5c)**. Of note, the frequencies of *FOXP3^+^* naïve and memory Treg cells in the blood of the patients with SjD were comparable to the controls.

**Figure 6.**
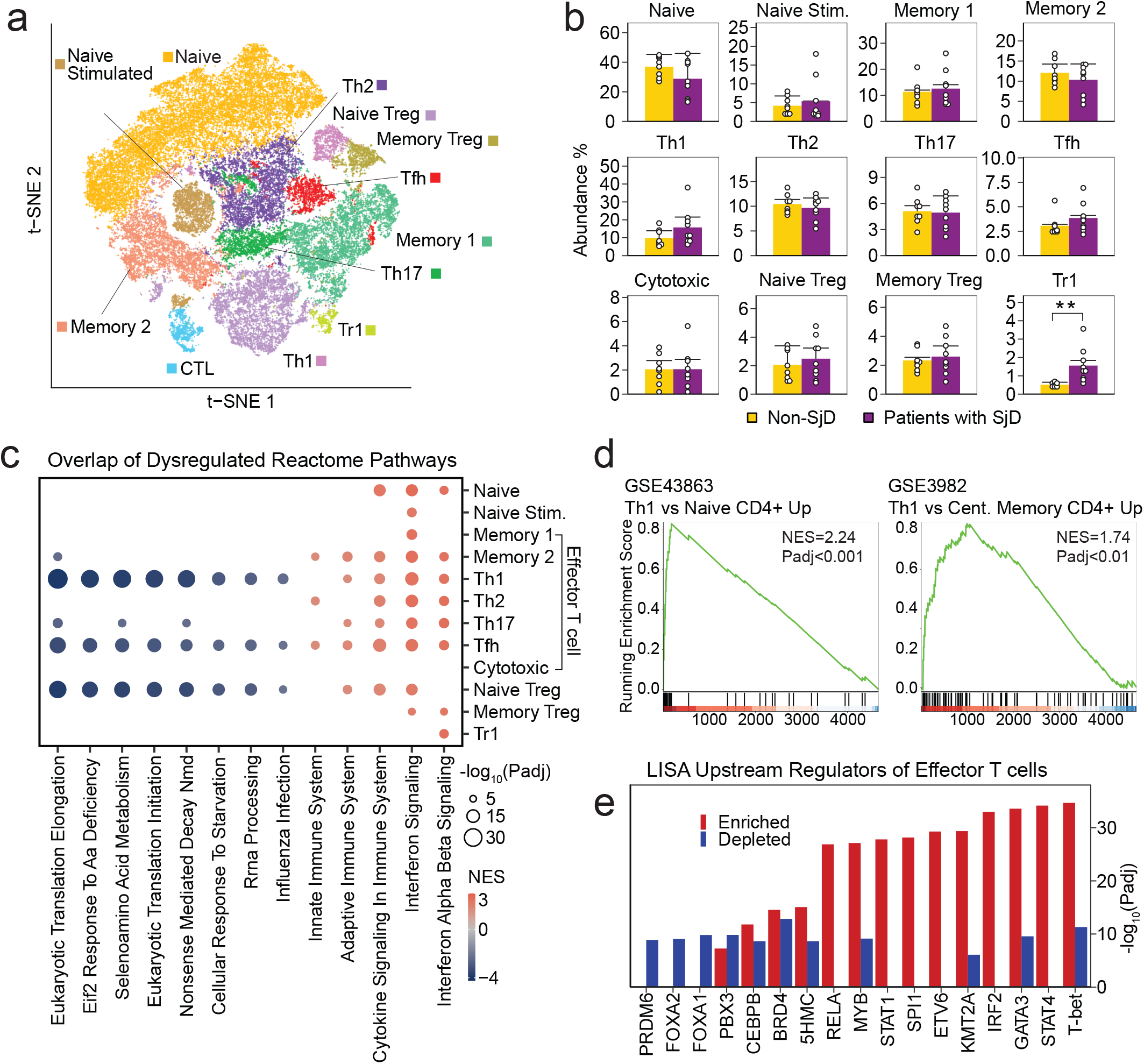
Increased IFN signaling in PBMCs of patients with SjD. **a)** Aggregate t-SNE reduction of 46,376 CD4^+^ T cells from human PBMCs of 9 patients with SjD and 8 symptomatic non-SjD controls. CD4^+^ T cell subsets were identified by marker gene expression. **b)** Frequencies of CD4^+^ T cell subsets in SjD and non-SjD control cohorts. **c)** Reactome pathway analysis of DEG within each CD4^+^ T cell subset of patients and controls. Pathways were considered significant if their *P* adj was < 0.01. Only pathways found to be dysregulated in more than two CD4^+^ T cell subsets were included in panel **c, d)** GSEA of effector CD4^+^ T cell subsets shown in panels **(a-c)** including memory, Th and cytotoxic T cells using the immunological signature database. **e)** Top 10 enriched and depleted upstream regulators of DEG in all effector CD4^+^ T cell subsets using LISA upstream regulator analysis. All statistical analyses were conducted using the Wilcoxon rank-sum test, and significance was adjusted using the Benjamini-Hochberg method. Results are shown as means ± SD. \**P* < 0.05, \*\**P* < 0.01, \*\*\**P* < 0.001.

To glean information about the activation status and function of CD4^+^ T cells in the patients with SjD, we analyzed the DEGs for each T cell subset and performed a functional enrichment test (FET) to determine if the DEGs were associated with specific pathways or cellular function. CD4^+^ T cell subsets shared similar patterns of dysregulated pathways, most prominently an enrichment of interferon (IFN) signaling **(Figure 6c)**. Both type I (IFN-α/β) and type II (IFN-γ) IFN signaling pathways were strongly and significantly enriched in the patients with SjD compared to the non-SjD controls. This finding is consistent with greater type I and II IFN signatures reported in patients with SjD^71,72^ and the Th1/IFN-γ dominated inflammation in SGs of *Stim1/2^Foxp3^* mice. Several CD4^+^ T cell subsets, including Th1, Tfh and naïve Treg cells, showed a depletion of ribosome-related pathways, which was also observed in other scRNA-seq studies of patients with SjD^73^. This is of note as auto-antibodies in SjD are directed against Ro and La ribonucleoproteins that are associated with the quality control of 5S ribosomal RNA^74^ and maturation of RNA polymerase III^75^, potentially explaining the dysregulation of ribosomal related pathways in some CD4^+^ T cell subsets.

To further understand the mechanisms by which CD4^+^ T cells cause inflammation in SjD, we used DEGs from effector CD4^+^ T cells, including Th1, Th2, Th17, Tfh, cytotoxic and memory T cells (but excluding naive and Treg subsets), to conduct in-depth pathway and upstream regulator analyses. We observed a significant enrichment of genes in pathways associated with oxidative phosphorylation, interferon signaling, Th1 and Th2 pathways in patients with SjD compared to controls **(Supplementary Figure 5d)**. An enrichment of Th1 cell-associated genes was also apparent by GSEA analysis using the immunological signature database **(Figure 6d)**. An upstream regulator analysis using LISA to predict transcription factors (TFs) regulating gene expression in effector CD4^+^ T cells of patients with SjD identified several TFs associated with Th1 and Th2 cell differentiation and function, including T-bet, STAT1, STAT4 and GATA3, respectively **(Figure 6e)**. Together, these data provide evidence for an enhanced Th1 response in effector CD4^+^ T cells of patients with SjD compared to non-SjD controls, which is similar to our findings in *Stim1/2^Foxp3^*mice.

Given the enhanced IFN signaling and Th1 gene expression and TF signatures we found by the analysis of all effector CD4^+^ T cell subsets, we investigated whether Th1 cells are hyperactivated. Using the Louvain algorithm-based clustering, we identified a distinct cluster of *IL-12RB^+^ STAT1^+^ STAT4^+^ IFNG^+^*Th1 cells **(Figure 7a)** on which we performed pathway analyses of DEGs. A pathway analysis using the IPA platform revealed an enrichment of interferon signaling and Th1-associated pathways **(Figure 7b)**, which was confirmed by GSEA analysis that showed an increase in pathways related to the positive regulation of cytokine and IFN-γ production **(Figure 7c)**. Of note, the eukaryotic initiation factor 2 (eIF2) signaling pathway was significantly downregulated in Th1 cells of patients with SjD **(Figure 7b)**. eIF2 is an essential factor for the initiation of protein synthesis, and depletion of eIF2-related gene expression in Th1 cells is consistent with the reduced eukaryotic translation initiation signatures we had observed in several CD4^+^ T cell subsets **(Figure 6c)**. To characterize functional states of CD4^+^ T cells in patients with SjD in an unbiased manner, we performed non-negative matrix factorization (NMF). NMF identifies common gene expression profiles that can reflect cell type identity, as well as their functional state^76^. We detected 23 gene expression programs (GEPs) in CD4^+^ T cells of patients with SjD and controls, including a Th1-specific GEP **(Figure 7d)** that mapped predominantly to the Th1 cell cluster we had identified by using Th1-specific marker genes **(Figure 7a)**. Importantly, the GEP associated with Th1 function was significantly enriched in Th1 cells of patients with SjD compared to the non-SjD controls **(Figure 7e).** This enrichment of a functional Th1 signature was most prominent in Th1 cells but could also be detected in other CD4^+^ T cell subsets, including CTLs and memory 1 and 2 cells **(Figure 7e)**. The majority of Th1 functional genes, such as *IFI27, IL12RB2, CD74* and *STOM*, were upregulated in the Th1 subset of patients with SJD compared to the controls **(Figure 7f)**. Collectively, these data demonstrate an enhanced function of Th1 cells, including IFN signaling, despite normal Th1 frequencies.

**Figure 7.**
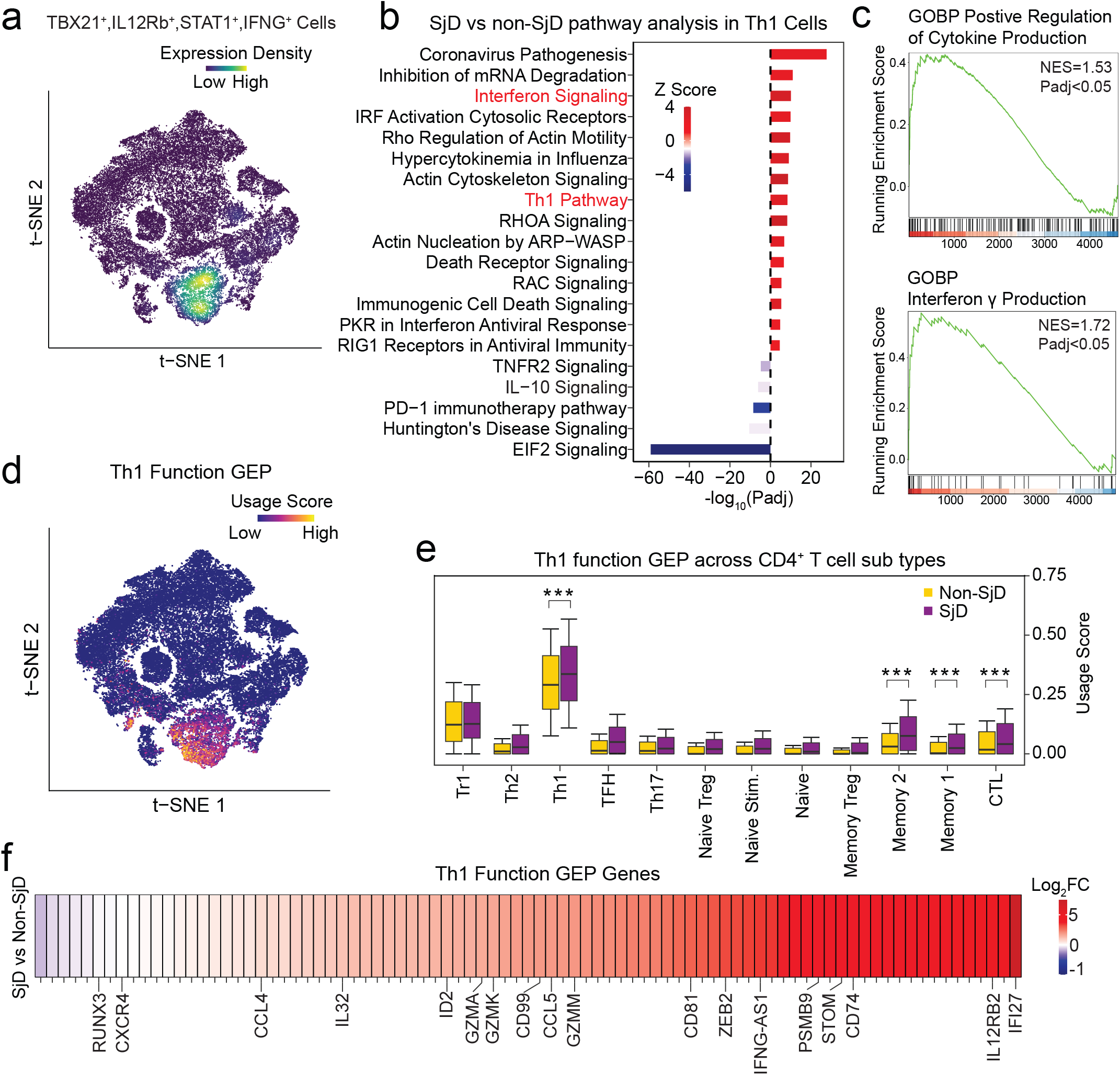
Th1 signature in PBMCs of patients with SjD. **a)** t-SNE representation of CD4^+^ T cells expressing Th1 marker genes (*IL12Rb, STAT1, STAT4, IFNG*) from 9 patients with SjD and 8 symptomatic non-SjD controls. **b)** Pathway enrichment analysis based on DEGs in Th1 cells using the IPA database. Top 20 dysregulated pathways ranked by *P* value (-log10). Z scores of pathway enrichment / depletion are indicated. **c)** GSEA of Th1 cells from SjD patients and controls using the immunological signature database. **d)** t-SNE representation of CD4^+^ T cells with a *Th1 Function* gene expression profile (GEP) identified by non-negative matrix factorization (NMF). **e)** Usage scores for *Th1 Function* GEP by CD4^+^ T cell subset. **f)** Tileplot of *Th1 Function* GEP genes that are differentially expressed in Th1 cells of patients with SjD compared to non-SjD controls. All statistical analyses were conducted using the Wilcoxon rank-sum test. Significance was adjusted using the Benjamini-Hochberg method. Results are shown as means ± SD. \**P* < 0.05, \*\**P* < 0.01 and \*\*\**P* < 0.001.

### Diminished function of memory Treg cells in patients with SjD

Deletion of *Stim1/2* abolishes SOCE and impairs Treg function^42^, resulting in Th1 cell-mediated severe SjD-like disease of *Stim1/2^Foxp3^* mice. These findings indicate that Treg cell function is required to prevent autoimmune inflammation of exocrine glands in mice. We therefore hypothesized that Treg function might also be perturbed in patients with SjD, although the role of Treg cells in the pathogenesis of SjD remains ambiguous. Most studies have focused on analyzing the frequencies of Treg cells in the blood or SGs of patients with SjD, which has yielded conflicting results^25-34^. We found that the frequencies of Foxp3^+^ naive and memory Treg cells were unchanged and those of Foxp3^-^ Tr1 cells greater in patients with SjD compared to non-SjD controls **(Figure 6b)**. To assess the function of Foxp3^+^ Treg cells, we combined naïve and memory Treg populations to have sufficient cell numbers for further analysis **(Figure 8a)**. While the capture rate by scRNA-seq in Tregs was not sufficient to analyze the transcript levels of most genes controlling Ca^2+^ homeostasis, including *STIM1* and *ORAI1* **(Supplementary Table 4)**, our analysis of all CD4^+^ T cells showed only moderate (albeit significant) changes in their mRNA levels **(Supplementary Figure 5e, f)**. More conclusive was an IPA upstream regulator analysis of Treg subsets that showed a significant enrichment of molecules involved in type I, II, and III IFN signaling, including *IFNA2*, *IFNG*, *IFNL1* and IRF7 **(Figure 8b)**. This finding was confirmed by GSEA of immunological signature databases that identified a significant enrichment of IFN signaling **(Figure 8c).** Foxp3^+^ Treg cells had been reported to upregulate T-bet in response to IFN-γ, resulting in Treg cells with unique properties optimized for the suppression of Th1 responses^77^. However, this is not the case here because the Th1 functional GEP we had identified earlier was enhanced in Th1 cells, memory and cytotoxic CD4^+^ T cells but not in the Treg populations of patients with SjD **(Figure 7e)**. Instead, the enhanced IFN signature of Treg cells likely results from increased IFN levels in SjD, which affects all CD4^+^ T cell subsets **(Figure 6c)**. Of note, a similar enrichment of several IFN response genes, such as *Stat1, Gbp1, Cxcl9*, was observed in Treg cells isolated from the cLN of male *Stim1/2^Foxp3^* mice **(Supplementary Figure 5g)**.

**Figure 8.**
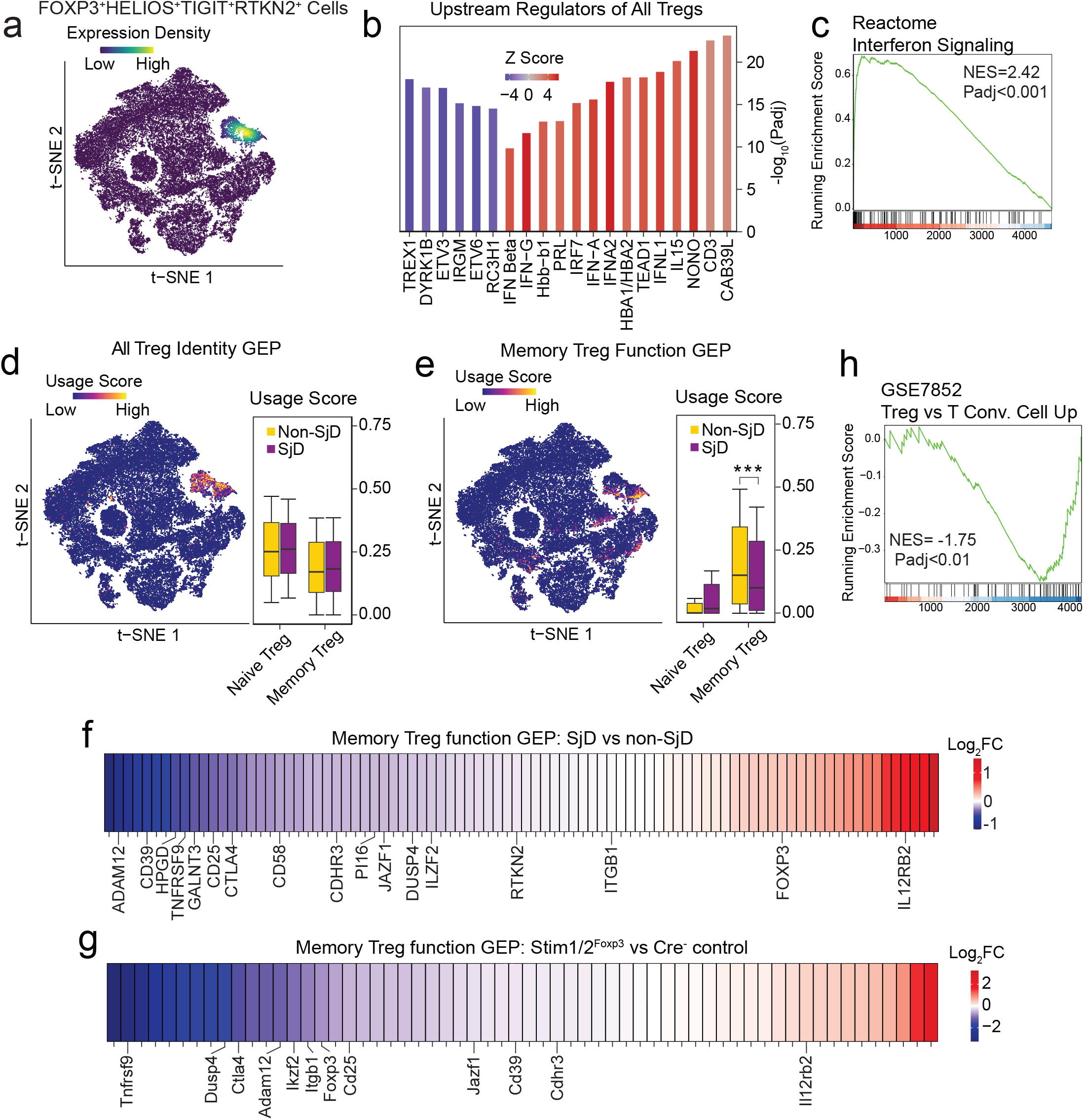
Attenuated functional memory Treg cell signature in PBMCs of patients with SjD. **a)** t-SNE representation of CD4^+^ T cells expressing Treg marker genes *(FOXP3, HELIOS, TIGIT, RTKN2)* from 9 patients with SjD and 8 symptomatic non-SjD controls. **b)** Top 20 depleted and enriched upstream regulators of DEG in all Treg cells using the IPA upstream regulator analysis. **c)** Selected GSEA of all Treg cells from patients with SjD and controls using the immunological signature database. **d)** t-SNE representation of CD4^+^ T cells with an *All Treg Identity* GEP identified by NMF (left) and usage scores of this GEP by naive and memory Treg cells of SjD patients and controls (right). **e)** t-SNE representation of CD4^+^ T cells with a *Memory Treg Function* GEP identified by NMF (left) and usage scores of this GEP by naive and memory Treg cells of SjD patients and controls (right). **f)** Tileplot of *Memory Treg Function* GEP genes that are differentially expressed in memory Treg cells of patients with SjD compared to non-SjD controls. **g)** Tileplot of *Memory Treg Function* GEP genes that are differentially expressed in Foxp3^+^ YFPcre-positive and YFPcre-negative control Treg cells from heterozygous female *Stim1/2^Foxp3^* mice^42^. **h)** Selected GSEA of all Treg cells from patients with SjD and controls using the immunological signature database. All statistical analyses were conducted using the Wilcoxon rank-sum test. Significance was adjusted using the Benjamini-Hochberg method. Results are shown as means ± SD. \**P* < 0.05, \*\**P* < 0.01 and \*\*\**P* < 0.001.

Alterations in Treg cell function have been reported in several autoimmune diseases, such as T1D, MS and RA^78-81^. To investigate if Treg cells are dysfunctional in patients with SjD, we analyzed scRNA-seq data of PBMC from SjD and non-SjD samples by NMF with the goal to determine GEPs representing Treg function. We identified 2 GEPs, of which one was an *All Treg identity* GEP expressed by both naïve and memory Treg populations (**Figure 8d)**. The other was a *Memory Treg function* GEP predominantly expressed by memory Treg cells (**Figure 8e)**. Whereas we did not detect quantitative changes in *All Treg identity* GEP usage between PBMCs of patients with SjD and non-SjD controls for the *All Treg identity* GEP **(Figure 8d),** the *Memory Treg function* GEP was significantly depleted in patients with SjD **(Figure 8e)**. By manual inspection of individual genes of the *Memory Treg function* GEP, we found that many genes associated with Treg function, such as *CD39, TNFRSF9, CD25 and CTLA4*, were depleted in PBMCs of patients with SjD, although *FOXP3* was not one of them **(Figure 8f)**. To determine if the depletion of the *Memory Treg function* GEP in PBMCs of patients with SjD correlated with the functional status of Treg cells in *Stim1/2^Foxp3^*mice, we assessed the expression of the mouse homologs of GEP genes in Treg cells isolated from *Stim1/2^Foxp3^* mice^42^. (Note that we used sorted *Stim1/2*-deficient YFP^+^ Treg cells from female heterozygous *Stim1/2^Foxp3^* mice that do not develop autoimmunity to avoid secondary effects on gene expression.) Although not all genes belonging to the human *Memory Treg GEP* had homologs in mice, 57 of 77 genes depleted in PBMCs of patients with SjD were also found to be reduced in *Stim1/2*-deficient mouse Treg cells **(Figure 8g)**, suggesting a significant correlation of Treg dysfunction in patients with SjD and *Stim1/2^Foxp3^* mice. To further confirm the memory Treg dysfunction in patients with SjD, we conducted a GSEA analysis on the memory Treg subset **(Figure 6a)** and found a significant depletion of Treg signature genes in patients with SjD compared to non-SjD controls **(Figure 8h)**. Collectively, these data indicate that antigen-experienced memory Treg cells are dysfunctional in patients with SjD and may thus, contribute to the patients’ exocrine gland autoimmunity.

## DISCUSSION

The causes of SjD remain inadequately understood. Animal models of SjD have uncovered important aspects of the pathophysiology of the human disease^82^, but no model reflects all clinical and immunological characteristics of human SjD^82-84^. The most common models are based on nonobese diabetic (NOD) mice and its derivatives, such as NOD.B10-H2^b^ and C57BL/6.NOD-Aec1Aec2 mice, which show immune cell infiltration of exocrine glands, and reduced SG and LG function^82,83^. Other SjD models are based on knockout mice in which the genes coding for Id3 (Inhibitor of differentiation 3), aromatase or phosphatidylinositol 3-kinase (PI3K) are deleted, and transgenic mice overexpressing B-cell-activating factor (BAFF) or lymphotoxin alpha (LTα). Some models induce SjD through viral infection or by immunization with potential autoantigens such as Ro, M3R, and carbonic anhydrase II (CAII)^82^. These models recapitulate some but not all features of human SjD and either lack lymphocytic SG and LG infiltration, gland dysfunction or generation of SjD-associated autoantibodies.

Here, we report that conditional deletion of *Stim1* and *Stim2* in FoxP3^+^ Treg cells results in a severe form of SjD-like disease in mice which has all the clinical features observed in humans with SjD^36^. Notably, *Stim1/2^Foxp3^* mice show severe lymphocytic infiltration of the SG and the LG resulting in fibrosis and destruction of the glands over time, SG and LG dysfunction with markedly reduced saliva and tear production, and elevated serum levels of anti-Ro/SSA and anti-La/SSB autoantibodies. We should note that we had shown in a previous study that Treg cells in *Stim1/2^Foxp3^* mice fail to differentiate into follicular Treg (Tfr) cells that control the germinal center (GC) reaction between Tfh cells and GC B cells^42^. This lack of Tfr cells very likely, then, explains the expansion of autoreactive B cells and the production of autoantibodies including those against anti-Ro/SSA and anti-La/SSB antigens.

Extraglandular disease manifestations are present in 50-60% of patients with SjD^1,2^, which often include pulmonary disease with either interstitial pneumonia or lymphocytic interstitial pneumonia (LIP)^49^. Pulmonary inflammation with perivascular and peribronchial immune cell infiltration were also present in *Stim1/2^Foxp3^* mice. All SjD-like symptoms were present in *Stim1/2^Foxp3^* mice by 4 weeks of age, which is notably shorter than in other SjD disease models^12,85-89^. A notable difference between human SjD cases and *Stim1/2^Foxp3^* mice is that SjD mostly affects women with a female-to-male ratio of ∼ 1:16-20^90,91^, whereas *Stim1/2^Foxp3^* mice are all male owing to the fact that the *Foxp3* gene driving Cre expression is located on the X chromosome. (Note that in female heterozygous female *Stim1/2^Foxp3^* mice half of all Treg cells are WT, and, thus, the female heterozygous mice consequently do not develop disease). Notwithstanding this sex difference, we propose that *Stim1/2^Foxp3^* mice are a valuable new, rapid and more accurate model of SjD that can help to shed novel insights into the pathogenesis of human disease because *Stim1/2^Foxp3^* mice not only have all the clinical manifestations of human SjD they also share many molecular disease features. Our comparison of transcriptional profiles in the SG from *Stim1/2^Foxp3^* and WT mice and SGs of patients with SjD and controls^44^ revealed that 70% of all DEGs selectively upregulated in the SGs of patients with SjD were also upregulated in the SG of *Stim1/2^Foxp3^* mice. Moreover, gene set enrichment and pathway analysis showed strong dysregulation of genes related to type 1 IFN production and IFNα/β signaling in the SG of *Stim1/2^Foxp3^* mice, which is reminiscent of the increased type 1 IFN levels and upregulated expression of type I IFN-stimulated genes found in exocrine glands and the blood of patients with SjD^60,61^. A similar overlap of dysregulated gene expression in patients with SjD and *Stim1/2^Foxp3^* mice is evident from the fact that ∼ 50% of differentially methylated genes identified in patients with SjD^43^ are differentially expressed in the SG of *Stim1/2^Foxp3^* mice. Together, our data demonstrate that SjD in patients and SjD-like disease in *Stim1/2^Foxp3^* mice share many phenotypic, as well as cellular and molecular immunological features, pointing to a shared disease pathophysiology.

Besides autoantibodies against Ro/SSA, La/SSB and other autoantigens, SjD is characterized by T cell infiltration of the SG and the LG^52^. The role of autoreactive T cells in exocrine gland inflammation and dysfunction was revealed in mouse models in which keratoconjunctivitis and LG inflammation induced by desiccating stress could be transferred to non-stressed nude mice by adoptive transfer of CD4^+^ T cells^9^. LG inflammation could also be induced in NOD-SCID mice by transfer of CD8^+^ T cells from NOD mice^10^. Moreover, T cells from mice immunized with M3R peptides were able to induce sialadenitis and cause hyposalivation after adoptive transfer to lymphopenic *Rag1^-/-^* mice^15^. These studies suggested that autoreactive T cells are main drivers of exocrine gland inflammation and dysfunction independent of B cells and autoantibodies. A study of different CD4^+^ T cell subsets present in the SG of patients with SjD found that ∼30% of CD4^+^ T cells expressed T-bet identifying them as Th1 cells compared to 20-30% Tfh cells, 10-20% CD4^+^ CTL, 5-10% Th2 cells, and 2-10% Th17 cells^65^. Moreover, the expression of T-bet, IFN-γ and TNF was found to be increased in the SG and saliva of patients with SjD^54-58,64,65^, and the expression of IFN-γ by lymphocytes isolated from the labial SG of patients with SjD correlated with the degree of lymphocytic infiltration^54^.

Here, we show that transcriptional changes in the inflamed SG of *Stim1/2^Foxp3^* mice are dominated by Th1 and Th2-related pathways. RT-PCR data, as well as pathway and upstream regulator analyses of RNA-seq data, showed an enrichment of Th1 and Th2 signature genes, such as IFN-γ, TNF and IL-4, and transcription factors that regulate Th1 and Th2 differentiation, including T-bet, STAT1, GATA3 and STAT6. These findings are consistent with altered Th1 and Th2 cytokine levels in patients with SjD^44,54-58,64-66^. We demonstrate the importance of Th1 cells for SjD-like disease in mice by showing that the adoptive transfer of CD4^+^ T cells from *Stim1/2^Foxp3^* mice to lymphopenic *Rag1^-/-^* mice, which lack B cells and antibody production, is sufficient to transfer SjD-like disease, including SG and LG inflammation and dysfunction. (It is noteworthy that transfer of CD8^+^ T cells from *Stim1/2^Foxp3^* mice did not induce disease.) The transferred CD4^+^ T cells isolated from inflamed SG, LG and draining LNs of host mice had enhanced IFN-γ production and T-bet expression, suggesting that they had differentiated into Th1 cells.

To address how Th1 cells from *Stim1/2^Foxp3^* mice cause SjD-like disease, we deleted IFN-γ as the signature cytokine produced by Th1 cells. *Ifng* deletion in CD4^+^ T cells of *Stim1/2^Foxp3^* mice before adoptive transfer compromised their ability to cause SG and LG inflammation and dysfunction in *Rag1^-/-^* host mice. Our findings are consistent with earlier reports that showed that targeted deletion of IFN-γ or the IFNGR in NOD mice prevents lymphocytic infiltration of SG, restores saliva production and suppresses the generation of autoantibodies^92^. Moreover, deletion of IFN-γ in *Il2ra* (encoding Cd25) knockout mice delays lymphocytic infiltration of the LG and preserves its function before 8 weeks of age^93^. Besides suppression of IFN-γ signaling, anti-TNF-α treatment in NOD mice also significantly alleviates SG inflammation and rescues saliva production, which is associated with reduced T-bet protein levels, suggesting that TNF-α production by Th1 cells contributes to SG pathology^94^. The mechanisms underlying the important role of IFN-γ in SjD pathogenesis are not fully understood. Our data demonstrate that treatment of SG acinar cells with IFN-g suppresses Ca^2+^ influx, which is required for the activation of the Ca^2+^-activated chloride channel ANO1 (or TMEM16A) and thus, chloride secretion and saliva production^95,96^. Protection from SjD-like disease following suppression of IFN-γ signaling in CD4^+^ T cells therefore is due, at least in part, to preventing SG dysfunction. Our findings provide a mechanistic explanation for the pathogenic function of IFN-γ-producing Th1 cells and their ability to cause SjD-like disease.

Importantly, the pathogenic role of Th1 cells is not limited to our mouse model. Our scRNA-seq analysis of patients with SjD and non-SjD controls identified a distinct cluster of *IL-12RB^+^ STAT1^+^ STAT4^+^ IFNG^+^* Th1 cells, whose frequency was not increased in patients with SjD compared to controls. Pathway analyses of differentially expressed genes, however, revealed an enrichment of pathways related to IFN signaling and Th1 differentiation in patients with SjD. Using non-negative matrix factorization as an independent method to define functional states of CD4^+^ T cells in patients with SjD, we identified a Th1-specific gene expression program that was significantly enriched in Th1 cells of patients with SjD. The majority of Th1 functional genes, such as *IFI27, IL12RB2, CD74* and *STOM*, were upregulated in patients with SjD, suggesting enhanced Th1 cell function. Our data provide evidence for an enhanced Th1 response in effector CD4^+^ T cells of patients with SjD, which is consistent with previous reports in patients with SjD^54-57,64,71,72^ and similar to our findings in *Stim1/2^Foxp3^*mice.

In *Stim1/2^Foxp3^* mice, the severe SjD-like phenotype results from Treg dysfunction despite normal numbers of Treg cells. Deletion of *Stim1* and *Stim2* abolishes SOCE and impairs Treg-mediated immunosuppression, as we had shown previously^42^. The striking phenotypic and immunological similarities between patients with SjD and *Stim1/2^Foxp3^* mice point to an important role of Treg cells in the pathophysiology of SjD. Indeed, several mouse models have linked impaired Treg development or function to SjD-like disease. *Il2^-/-^* and *Il2ra^-/-^* mice that lack Foxp3^+^ Treg cells show lymphocyte infiltration in exocrine glands and hyposalivation^35^. Intriguingly, scurfy mice, which lack functional Treg cells due to a missense mutation in the *Foxp3* gene, do not develop a SjD-like phenotype unless they are fed with LPS or lymphocytes from scurfy mice are transferred to *Rag1^-/-^* hosts^35^. The lack of spontaneous SjD in scurfy mice despite the absence of Treg cells is most likely explained by their very short median survival time of 3-4 weeks, which is too brief to develop disease. *Id3^−/−^* mice, which have a defect in the TGF-β1-induced generation of thymus-derived Treg cells and enhanced abnormal Th17 responses^97^, develop a SjD-like disease, including dry eyes and mouth, lymphocyte infiltration in the SG and the LG and generation of anti-Ro and anti-La antibodies^12^. In several mouse models of SjD, such as NFS/sld, IQI/Jic, and Ar KO mice, thymectomy shortly after birth exacerbated disease symptoms, suggesting a protective role of thymus-derived Treg cells^82^. Aged NOD.B10.H2b mice, which spontaneously develop SjD-like disease, including LG gland inflammation, accumulate dysfunctional Treg cells with lower suppressive capacity^98^.

A role of Treg cells in human SjD has been suggested but the supporting data are ambiguous^99,100^. Most studies have focused on the abundance of Treg cells. Treg numbers in the MSG of patients with SjD were reported to be increased^30,32^ or reduced^25^. The former two studies reported a positive correlation between increased Treg frequencies in MSGs and lymphocytic infiltration (focus scores)^30,32^. In the peripheral blood, Treg frequencies were reported to be either unchanged, increased, or decreased^25-34^. The cause of these discrepancies is unclear, but it may be related to how Treg cells were defined (as CD4^+^CD25^+^ or CD4^+^Foxp3^+^ T cells) or due to analysis at different stages of SjD. More relevant for SjD pathophysiology than mere Treg numbers may be their actual function. One study reported a lower suppressive capability of CD4^+^CD25^+^ Treg cells in the blood of patients with SjD compared to healthy individuals^28^, whereas two others found normal function of Tregs from patients with SjD ^25,27^. In recent years, scRNA-seq has greatly enhanced the ability to characterize immune cell subsets in autoimmune diseases such as SLE, MS and RA, in terms of their frequency and functional states^101,102^. In SLE, one of the most common causes for secondary SjD^103,104^, scRNA-seq and ATAC-seq (assay for transposase-accessible chromatin with sequencing) of peripheral CD4^+^ T cells identified an increased frequency of Treg cells that were exhausted based on increased expression and chromatin accessibility of T cell exhaustion markers and reduced suppressive function *in vitro*^105^. Similar transcriptomic analyses of Treg cells in SjD are not yet available. One scRNA-seq-based study investigating 3 patients with SjD and 3 controls concluded that there are no changes in Treg frequencies in patients^73^, whereas another scRNA-seq study of 5 patients and controls did not mention Treg cells^66^. Here, we report that the numbers of Foxp3^+^ naive and memory Treg cells are comparable in 9 patients with SjD and 8 relevant controls. The frequencies of Foxp3^-^ Tr1 cells, by contrast are significantly greater in patients than controls, which is similar to an earlier report on patients with SjD^28^, but different from other autoimmune diseases, including psoriasis and MS, in which the frequencies of Tr1 cells were lower than in controls^106,107^.

To identify potential functional defects in Treg cells, we analyzed the transcriptomes of CD4^+^ T cells and Treg cells. We first looked for evidence of potential defects in Ca^2+^ signaling in patients with SjD because abolishing SOCE in Treg cells of *Stim1/2^Foxp3^* mice results in SjD-like disease and because several studies have previously reported dysregulated Ca^2+^ signaling in SjD. STIM1 and STIM2 protein levels were decreased in PBMCs of patients with SjD^108^ and Ca^2+^ influx was reduced in SG acinar cells from patients with SjD in response to carbachol stimulation, which was attributed to reduced expression of IP3R2 and IP3R3^109^. We used scRNA-seq data of patients with SjD to analyze transcript levels of regulators of Ca^2+^ homeostasis in T cells, which showed only moderate (albeit significant) changes in all CD4^+^ T cells (note that the scRNA-seq capture rate in Treg subsets was not sufficient to analyze transcript levels). While our analyses did not provide strong evidence for dysregulated Ca^2+^ signaling, at least at the transcriptional level, NMF analysis of scRNA-seq data revealed the depletion of a GEP associated with *Memory Treg function.* Depleted genes in this GEP included many that are critical for Treg function, such as CD25, CD39, CTLA4 and CD137 (4-1BB). Intriguingly, most genes in this GEP that were depleted in patients with SjD were also reduced in Treg cells of *Stim1/2^Foxp3^* mice. In addition, we found an increased IFN signature in all Treg cell subsets, which likely results from increased IFN levels in patients with SjD rather than indicating a Th1-like phenotype that had been identified in several human autoimmune diseases, including T1D, MS and RA^78-80^, and in a mouse model of SjD^81^. The IFN signature was present in all CD4^+^ T cell subsets of patients with SjD and not specific to Treg cells, and the Th1 functional GEP we identified in Th1 cells by NMF was not enriched in Treg cells of patients with SjD. Taken together, our findings show that the transcription of critical regulators of memory Treg is reduced in Foxp3^+^ Treg cells in patients with SjD. As these changes resemble those in Treg cells of *Stim1/2^Foxp3^* mice, which develop several aspects of SjD, we propose that impaired Treg function is a critical contributing factor to SjD in mice and human patients.

## KEY MESSAGES

### What is already known on this topic

- SjD is a common autoimmune disorder, but its etiology and pathophysiology are incompletely understood. Autoreactive B cells and autoantibodies are considered to be crucial for the pathogenesis of SjD, and have become a target for therapy, whereas the role of effector T cells and T regulatory (Treg) cells in SjD remains ambiguous.

### What this study adds

- *Stim1/2^Foxp3^*mice represent a new, more accurate and faster animal model of SjD that matches all diagnostic criteria of human SjD and recapitulates some of its extraglandular manifestations.
- *Stim1/2^Foxp3^*mice have normal Treg numbers, but the suppressive function of Treg cells is impaired, demonstrating that changes in Treg cell function, rather than merely their numbers, play an important role in preventing exocrine gland autoimmunity in SjD.
- Inflammation in salivary glands (SGs) of *Stim1/2^Foxp3^* mice is dominated by Th1 immune responses. Autoreactive, IFN-γ producing CD4^+^ T cells are sufficient to induce SjD-like disease in the absence of B cells and autoantibodies, suggesting the disease is primarily driven by this immune cell type rather than B cells.
- We identify increased Th1 responses and attenuated memory Treg function in CD4^+^ T cells of patients with SjD, which is consistent with the immune dysregulation observed in *Stim1/2^Foxp3^* mice.

### How this study might affect research, practice or policy

- Our study demonstrates an important role of Treg cells in SjD pathogenesis and stipulates a renewed attention to Treg cells in human SjD using single-cell transcriptomics and other omics approaches.
- The critical role of autoreactive, IFN-γ producing CD4^+^ T cells offers new insights into the pathogenesis of SjD and argues in favor of developing therapeutic strategies that target CD4^+^ T cells and in particular IFN-γ signaling to treat SjD in addition to B cell-centric therapeutic approaches.

## MATERIALS AND METHODS

Detailed experimental procedures, analyses and reagents are provided in Supplementary Materials.

### Acknowledgements

Patient data and specimens used in this manuscript are from the Sjögren’s International Collaborative Clinical Alliance (SICCA) Next Generation Studies, funded under grant #1U01DE02889 (and previously under contracts N01DE-32636 and #HHSN26S201300057C) by the National Institute of Dental and Craniofacial Research (NIDCR). This manuscript was prepared using a publicly available SICCA data set and does not necessarily reflect the opinions or views of the SICCA investigators, the NIH or NIDCR. We thank Dr. Jay Chiorini (NIDCR) for sharing the NS-SV-TT-AC salivary gland cell line.

### Contributors

Y.W., W.L., G.S., G.M., B.S., and M.V. conducted experiments. Y.W., W.L., M.M., G.S., G.M., F.Z., A.T., D.R., A.L.M., M.V., B.N., S.C., R.S.L. and S.F. analyzed data and interpreted the results. Y.W., W.L., G.S., G.M., M.V., S.C., R.S.L. and S.F. designed the experiments. Y.W., W.L., M.M., and S.F. wrote the manuscript. All authors read and approved the final version of the manuscript. Y.W. and W.L. contributed equally to this work.

### Funding

This study was funded by National Institutes of Health (NIH) grant R01DE027981 (to R.S.L. and S.F.), NIH grant EY030917 (to S.C.), U01DE028891 (to Caroline H. Shiboski, SICCA), NIH F30 training grant AI164803 (to A.Y.T.), an award from the Colton Center for Autoimmunity at NYU (to S.F.), and Hunan Province Graduate Student Research and Innovation Project CX20190160 from Central South University, Changsha, Hunan, China (to W.L.).

### Patient and public involvement

Patients and/or the public were not involved in the design, or conduct, or reporting, or dissemination plans of this research.

### Patient consent for publication

Not applicable.

### Ethics approval

Animal studies were conducted in accordance with the institutional guidelines for animal welfare approved by the Institutional Animal Care and Use Committee (IACUC protocol IA16-01359) at NYU Grossman School of Medicine. This study involves human participants and was approved by the University of California, San Francisco IRB committee (IRB# 10-02551). Participants gave informed consent to participate in the study before taking part in the study

### Data and materials availability

All data needed to evaluate the conclusions in the paper are present in the paper and/or the Supplementary Materials. Bulk and scRNA-Seq data have been deposited in the GEO database under accession numbers GSE253188 and GSE253568.

