## Supplementary materials for "Regulatory T cells and IFN-γ-producing Th1 cells play a critical role in the pathogenesis of Sjögren’s Syndrome"

### MATERIALS AND METHODS

**Mice.** *Stim1*$^{fl/fl}$*Stim2*$^{fl/fl}$ have been described previously[1,2] and were crossed to *Foxp3-YFP-Cre (B6.129(Cg)* $^{Foxp3tm4(YFP/icre)Ayr}$*/J*) mice (The Jackson Laboratory, JAX, strain 016959). *Rag1*$^{-/-}$ mice *(B6.129S7-Rag1*$^{tm1Mom}$*/J)* were purchased from JAX (strain 002216). All mice were maintained on a *C57BL/6* genetic background and housed under SPF conditions. Genotyping was performed by Transnetyx (Cordova, TN). Male and female mice between 5 and 24 weeks of age were used. All experiments were conducted in accordance with the institutional guidelines for animal welfare approved by the Institutional Animal Care and Use Committee (IACUC) at NYU Grossman School of Medicine. Mice received standard rodent chow and autoclaved drinking water *ad libitum*.

**Patient samples.** Patient data and peripheral blood mononuclear cell (PBMC) samples of patients with SjD and Non-SjD controls were acquired from the Sjögren's International Collaborative Clinical Alliance (SICCA)[3,4]. The age, gender, autoantibody status, focus score and the 2016 ACR-EULAR SjD classification criteria score are shown in **Supplementary Table 3**.

**Cell Culture**. For isolating primary mouse acini, submandibular glands (SMGs) were excised, and connective and fat tissues were removed. SMGs were finely minced and digested in Dulbecco's modified eagle medium (DMEM, Gibco, 11960-044) containing 0.25 mg/ml of Liberase (TL, Roche, 05401020001) and 1% BSA (Sigma, A7888) for 25 min. The digested tissues were spun down (2000 rpm, 5 min) and cell pellets were washed and re-suspended in DMEM with 1% BSA. The digestion medium and SMG cells were continuously gassed with a mixture of 95 % $O_2$ and 5% $CO_2$, and incubated in a shaker/water bath at 37°C. The human submandibular acinar cell line (NS-SV-TT-AC) was obtained from Dr. John A. Chiorini at NIDCR/NIH. Cells were cultured in Keratinocyte SFM (K-SFM, Gibco, 10744019). Cells were harvested by treating them with 0.05 % Trypsin/EDTA (Gibco, 25300054) and subcultured about 3 x $10^5$ cells in 100 mm dishes at 37°C in a humidified 20% $O_2$, 5% $CO_2$ incubator for 4-5 days.

**Intracellular $Ca^{2+}$ measurements.** $Ca^{2+}$ measurements in mouse Treg cells were described previously[14]. Briefly, FACS-sorted Treg cells were labelled with 2 mM Fura-2-AM (Life Technologies; F1201) for 30 min in RPMI medium at room temperature and attached for 10 min to 96-well imaging plates (Fisher, Cat. 08-772-225) coated with 0.01% poly-L-lysine (w/v) (Sigma-Aldrich, P8920) and washed two times with $Ca^{2+}$-free Ringer solution (155 mM NaCl, 4.5 mM KCl, 2 mM $CaCl_2$, 1 mM $MgCl_2$, 10 mM D-glucose and 5mM Na-HEPES). Changes in intracellular $Ca^{2+}$ concentration were analyzed using a FlexStation 3 plate reader (Molecular Devices) at 340 and 380nm excitation wavelengths. Treg cells were stimulated with 1 µM

Thapsigargin (EMD Millipore, 586005) in $Ca^{2+}$-free Ringer solution and SOCE was analyzed after re-addition of 1mM $Ca^{2+}$ (final) Ringer solution. SOCE was quantified by the slope and peak of the F340/380 ratio, ER calcium store depletion was analyzed by the integrated $Ca^{2+}$ signal (area under the curve, AUC) after Thapsigargin stimulation and before addition of extracellular $Ca^{2+}$.

For measurements of cytosolic $Ca^{2+}$ concentrations ($[Ca^{2+}]_{cyt}$) in acinar cells, cells were loaded with 10 µM Fura-2-AM (Invitrogen, F1221) in DMEM with 1 % BSA, and incubated at 37 °C for 45 min. Cells were continuously gassed with a mixture of 95 % $O_2$ and 5 % $CO_2$. Cells were washed 3 times with normal Ringer's solution (pH 7.4) as follows (mM): 155.0 NaCl; 4.5 KCl; 2 $CaCl_2$; 1 $MgCl_2$; 10 D-glucose; and 10 HEPES (Sigma-Aldrich, USA), and kept on ice. Cells were allowed to attach to glass cover slips coated with Cell-Tak (Corning, 354240) for 10 min, and then incubated with 1 µM thapsigargin for 10 min at RT. During measurements of $[Ca^{2+}]_{cyt}$, cells were continuously perfused with Ringer's solution using a perfusion system (VC-6/8 valve controller) at 4 ml/min which was controlled by electrical control valves (Harvard Bioscience Inc., USA). Measurements were recorded first in $Ca^{2+}$- free Ringer's solution for 1 min followed by readdition of standard Ringer's solution for 3 min at room temperature. Fura-2 fluorescence was measured using a Nikon Ti2-E Eclipse microscope (Nikon) with an objective Nikon S Fluor x20 and a DS-Qi2 digital SLR camera (Nikon) controlled by NIS elements software (Version 5.20.01, Nikon). Cells were excited at 340 and 380 nm with an emission of 510 nm. Images were acquired every 5 sec and 340/380 ratio was calculated.

For measurements of $[Ca^{2+}]_{cyt}$ in human submandibular acinar cells (NS-SV-TT-AC), cells were plated on glass coverslips coated with laminin (Corning, 354239 and loaded with 1 µM of Fura2-AM for 30 minutes at room temperature in the dark and washed in normal Ringer's solution (pH 7.4). Cells were stimulated with either ATP (100 µM) or Carbachol (CCh; 1000 µM) in normal Ringer's solution. During $[Ca^{2+}]_{cyt}$ measurements, cells were continuously perfused with the indicated solution by a perfusion system at 4 ml/min. Cells were excited at 340 and 380 nm, and fluorescence emission was measured at 510 nm. Images were taken every 3 sec and 340/380 ratio calculated and graphed using Prism software (Graph Pad).

**Tear Production and Salivary Flow Measurements.** Mice were anesthetized with intraperitoneal injection of a cocktail of ketamine (10 mg/mL) and xylazine (0.2 mg /mL) at 10 µL/g of body weight. Tear production was analyzed using the phenol red thread test (PRTT)[5]. The "Zone Quick" phenol red thread (Ayumi Pharmaceutical Corporation) was applied to the lateral corner of the mouse cornea with forceps and kept stationary. After 1 minute, the color change of the thread from white to red was measured (in millimeters) as an equivalent of lacrimal flow. The measurement was conducted sequentially, with the right eye measured first, followed by the left eye. Following tear production measurements, while the mice remained immobilized, subcutaneous injections of 0.5 mg/kg pilocarpine (Sigma, P6503) were administered around the necks of mice. Subsequently, saliva secretion was monitored for a 20-minute duration by placing filter paper under the tongues of mice, and saliva production was quantified by measuring the weight change of the filter paper plus collection tube.

**Corneal Fluorescein Staining.** Mice were anesthetized with an intraperitoneal injection of ketamine and xylazine. To visualize the corneal damage, 10 µl of 0.125% Fluorescein sodium benoxinate hydrochloride ophthalmic solution (Altaire Pharmaceuticals, Inc) was added to each eye for 1 min and washed with sterile saline to eliminate excess dye and minimize background fluorescence. Eyes were wiped with sterile cotton Q tips and imaged on a Nikon SMZ 1500 dissection microscope equipped with Nikon Intensilight light source and a DAPI filter. Ocular surface damage was quantified by measuring the FITC intensity within the marked region of eye using ImageJ/Fiji software (NIH).

**Histology.** Tissue samples were fixed in 4% paraformaldehyde (Santa Cruz Biotechnology, Cat. SC-281692) for at least 24 hours, followed by a 30-minute wash in PBS, and two 30-minute washes in 70% ethanol. After the final wash, samples were immersed in 70% ethanol for 12 hours and subsequently embedded in paraffin before sectioning for H&E staining. For hematoxylin and eosin (H&E) staining of organs, including the SMG, the parotid gland (PG), the lacrimal gland (LG) and the lung, tissue samples were cut into 5 µm sections and stained with H&E using standard methods. Images were captured using a SCN400 slide scanner (Leica) and viewed through OMERO (Glencoe Software Inc.). For SMG and PG sections, a modified focus score was determined by a pathologist blinded to the experimental conditions by counting foci with more than 10 lymphocytes and calculating the number of foci per 4 $mm^2$ in whole gland sections. A focus score of 12 was given to cases in which inflammation was confluent. For lung sections, a pathologist blinded to the experimental conditions provided semi-quantitative assessment (mild/moderate/severe) of interstitial inflammation as well as binary assessment (presence/absence) of peribronchiolar and perivascular inflammation. The designation of “follicular bronchiolitis” was given to cases in which lymphoid follicles were identified in peribronchiolar inflammation, and the designation of “lymphocytic interstitial pneumonia” was defined by chronic bronchitis and bronchiolitis with dense lymphoid aggregates and lymphocytic infiltrate in the alveolar walls[6]. For image analysis of ocular histology, the degree of immune cell infiltration was quantified as the ratio of infiltrated ocular areas to the total area. Specifically, a random forest pixel classifier called iLastic[7] was used to identify basophilic pixels, which are presumed to belong to immune cells. Cuboid cells of the corneal epithelium, which are also basophilic, were manually excluded. The classifier was also used to identify eosinophilic pixels belonging to conjunctival connective tissue. The area of the basophilic pixels to the sum of basophilic and eosinophilic pixels within a region of interest (ROI) represent the level of immune cell infiltration. Calculations of infiltration, selection of ROIs, and exclusion of epithelial cells were performed with MATLAB/2022 (Mathworks).

**Quantitative Real-Time PCR**. Total RNA from SG lysates was extracted with the RNeasy Mini Kit (Qiagen, Cat. 74004) followed by cDNA synthesis using the iScript cDNA synthesis kit (BioRad, Cat. 1708891) according to the manufacturer's instructions. Quantitative Realtime (qRT-) PCR was performed using the Maxima SYBR Green qPCR Master Mix (Thermo, K0221) and a CFX96 Touch qRT Cycler (BioRad). Transcripts levels of target genes were quantified and normalized to the expression of housekeeping genes using the $2^{-\Delta CT}$ method. The primer sequences were listed in **Supplementary Table 1**.

**ELISA**. Ro52, Ro60 and La48 antigens (prepared as recombinant full-length protein as described[8]) were dissolved in PBS, added into a 96 well plate at 1 ug/well and incubated for at least 16 hours at 4°C. Wells were washed with solution of PBS with 0.05% Tween. Nonspecific binding sites were blocked with 0.1% gelatin/PBS. Mouse sera were diluted 1:50 in blocking solution and incubated for 1 hour on the plates. The plate was washed three times with PBS/Tween. Alkaline Phosphatase-conjugated rabbit anti-mouse IgG antibodies were added (at 1:3000, Sigma, Cat. AP160A) for 1 hour. Wells were washed with PBS/Tween and p-Nitrophenyl Phosphate, the substrate of Alkaline Phosphatase was added. Reactivity was monitored by OD at 405 nm using Spectramax (Molecular Devices) microplate reader and analyzed using Softmax Pro 7.1 software (Molecular Devices). Reactivity is reported in terms of Optical Density (OD)

**Bulk RNA-sequencing of mouse SMGs.** SMGs from 5-week-old Cre-negative littermate control and *Stim1/2*$^{Foxp3}$ male mice were harvested and kept in ice cold tubes with prefilled beads (Fisher, Cat. 15-340-153) in 1 mL TRIzol™ reagent (Invitrogen, 15596026). Organs were completely disassociated on tissue homogenizer (Bertin technologies, Precellys-24) at 6800 rpm for 30s. Total RNA was extracted as previously described[9]. RNA quality was assessed using a Bioanalyzer 2100 (Agilent) equipped with NANO chips. RNA libraries were prepared using automated stranded RNA-seq library prep polyA selection system (Illumina, TruSeq RNA Prep Kit). The libraries were purified with AMPure beads (Beckman Coulter), quantified using a Qubit 2.0 fluorometer (Life Technologies) and tested with an Agilent Tapestation 2200. RNA libraries were prepared using automated stranded RNA-seq library prep polyA selection system (Illumina, TruSeq RNA Prep Kit). The libraries were purified with AMPure beads (Beckman Coulter), quantified using a Qubit 2.0 fluorometer (Life Technologies) and tested with an Agilent Tapestation 2200. Prepared Libraries from different samples were pooled equimolarly and sequenced on a NovaSeq6000 system (Illumina) using a SP 100 Cycle Flow Cell v1.5 to generate 100 base pair paired-end reads. FASTQ files were generated using the bcl2fastq2 Conversion software for each sample (v2.20). Raw fastq files were assembled using Seq-N-Slide pipeline (10.5281/zenodo.5550459). Adaptors and quality control reads trimming was completed using trimmomatic v 0.36. FastQC was used to confirm quality of reads both before and after trimming. Trimmed paired end reads were then aligned using STAR v 2.7.7, with MM10 as the reference genome. The resulting binary alignment map files were converted into a gene count matrix using Feature count function of Bed tools v 2.10. Differential expression testing was completed using Bioconductor package DESEQ2 (v1.34), in an R statistical environment (v4.1.1). Genes with lower expression than 10 raw counts across in half of the total number of samples were filtered. Genes were considered significant if the adjusted p value was ≤ 0.01. Log fold shrinkage was calculated using native APEGM function of DESEQ2. The RNA-seq has been deposited in the GEO database under accession number GSE253188.

**Pathway Analysis**. Differentially expressed genes were utilized to perform a Functional Enrichment Test (FET) using the GSEA function of ClusterprofileR 4.0[10]. Databases included the canonical databases of the Molecular Signature Database, all three gene ontologies, and the Hallmark GSEA pathways datasets[11]. Additionally, immunological gene sets from the Molecular Signature Database were used to assess known functional differences between cell types. Pathways were considered significant if they had a Bonferroni adjusted value ≤ 0.01. GSEA plots were made using Gseaplot2 function of ClusterprofileR. The enrichment maps were calculated using the pairwise function in ClusterprofileR, and this similarity matrix was visualized in ggraph (v2.1.0). Edges between pathways not meeting a 30% shared gene threshold were removed. The Upstream regulatory analysis was generated using QIAGEN IPA (QIAGEN Inc). This method was applied to both our bulk RNA-seq and single-cell differentially expressed genes.

**Immunofluorescence.** Excised SMGs were fixed in 4% of paraformaldehyde (PFA) in phosphate-buffered saline (PBS) (Thermo scientific, Cat. J19943-K2) and embedded in paraffin and sectioned ~5 μm thick. For IF, unstained tissue sections were de-paraffinized using xylene (Fisher scientific, Cat. UN1307) followed by rehydration with different % of ethanol (From 100% to 70%, Pharmco, Cat. 111000190) and the antigen retrieval step with citric buffer (pH 6, Sigma, Cat. C9999) in heat for 20 min. Sections were then blocked with 5 % of skim milk (Bio-Rad, Cat. 170-6404) in PBS-T (Containing 0.1% of Tween-20) for 45 min at room temperature (22-25 °C) before applying primary antibody AQP5 (1:250, Rabbit, Ab92320, Abcam) in 5 % of skim milk in PBS-T overnight (4°C). After washing in PBS-T, secondary antibody Alexa flour 546 goat anti-rabbit IgG (H+L) (1:250, A11071, Life technologies) in 5% of skim milk in PBS-T was applied for 1 hour at room temperature. Sections were washed and mounted using Prolong gold antifade reagent with DAPI (Invitrogen, Cat. P36931). Fluorescence was measured using a Leica SP8 confocal microscope (Leica Microsystems) with objectives x10, x20, x40 (Oil) controlled by LAS X imaging software (Leica Microsystems). The acquired images were analyzed using ImageJ software (National Institutes of

Health).

**Flow Cytometry and Cell Sorting.** Lymph nodes were collected in 5mL cold PBS then ground through a cell strainer (70µm) (BD Falcon, Cat. 352350) using 20mL FACS buffer (2% FBS, 2mM EDTA in PBS). Cells were then centrifuged at 800g for 5 min. Cell pellet was resuspended in FACS buffer before staining. Salivary glands were collected in 5mL cold PBS, then minced into small pieces and incubated in 2mL "digestion mixture" with gentle shake at 50 RPM (1mg/mL collagenase D, 20 µg/ml DNase I, 10 mM HEPES, DMEM) for 45 minutes at 37°C in incubator. Digested tissues were ground up and cells were passed through a cell strainer (70µm). Cell strainer was rinsed with PBS and cells were centrifuged for 5 minutes at 800 x *g*. Cell pellet was resuspended as single cell suspension in FACS buffer for flow cytometry analysis. After blocking of Fc receptors using an anti-CD16/32 antibody (BioXcell, Cat. BE0307) and staining of dead cells using LIVE/DEAD Fixable Blue Dead Cell Stain Kit (ThermoFisher, Cat. L23105), cell surface molecules were stained at 4°C in the dark for 20 min at 4°C with the indicated antibodies. For transcription factor staining, surface-stained and LIVE/DEAD labeled cells were fixed with the Foxp3 / Transcription Factor Staining Buffer Set (eBioscience, Cat. 00-5523-00) for 30 min at RT in the dark and stained intracellularly at 4°C in 1x permeabilization buffer for 45 min. Acquisition of cells was performed on a FACS LSR-Fortessa flow cytometer (BD biosciences) and analyzed using the FlowJo 10.8.1 (TreeStar). Sterile cell sorting was performed on Aria-II cell sorter (BD biosciences). List of antibodies were shown in **Supplementary Table 2**.

**Adoptive T cell transfer.** Cervical lymph nodes from 4 to 5-week-old Cre-negative littermate control and *Stim1/2*$^{Foxp3}$ male mice were harvested and grounded through a 70 µm cell strainer using 10 mL sterilized PBS plus 2% FBS. Cells were stained with anti-CD4 (APC-Cy7), anti-CD25 APC), anti-CD8a (PE-Cy7) antibodies. Conventional CD4$^+$ T cells (CD4$^+$CD25$^-$Foxp3$^-$YFP$^-$) and CD8$^+$ T cells (CD8$^+$CD4$^-$) were sorted on Aria II (BD) followed by retroorbital injection of 1x10$^6$ cells into *Rag1*$^{-/-}$ recipient mice. Four weeks later, mice were sacrificed for gland function assay and histological analysis. For retrovirally transduced T cell transfer condition, CD4$^+$ T cells were first purified from cervical lymph node from *Stim1/2*$^{Foxp3}$ male mice by EasySep™ Mouse CD4$^+$ T Cell Isolation Kit (StemCell, Cat. 19852) with manufactural instruction. CD4$^+$ T cells were activated with anti-CD3e (20ng/mL) anti-CD28 (1 µg/mL) on 20 µg/mL rabbit-anti-hamster IgG (Thermo Scientific, Cat. A18891) coated 12-well plate with seeding density 1 x 10$^6$/mL. Twenty-four hours post activation, T cells were transduced by spin-transduction (2.500 rpm, 90 min, 32 °C) in the presence of retroviral supernatant and 8 µg/ml Polybrene (Sigma-Aldrich, Cat. 107689). Retroviral supernatant from T cells was removed 30 min after spin infection and replaced with fresh complete RPMI media. T cells were detached 24 h later and split into 6-well plate with 5mL of full RPMI media containing recombinant human IL-2 (20 U/mL) (PeproTech, Cat. 200-02) to expand for 3 more days before cell sorting. Retroviral supernatant was produced in the Platinum-E retroviral packaging cell line[12]. Platinum-E cells were transfected by GeneJet (Fisher, Cat. FERK0481) with retroviral expression plasmids *Lmpd-shCd19-pGK-Ametrine* and *Lmpd-shIfng-pGK-Ametrine* and the ecotropic packaging vector pCL-Eco. Retroviral supernatant was collected 36 and 60h after transfection. *shCd19* (control) and *shIfng* transduced $CD4^+$ T cells ($Ametrine^+$) were FACS-sorted using a sterile BD FACS AriaII cell sorter (BD Bioscience), and subsequently transferred into *Rag1*$^{-/-}$ host mice. Ten to twelve weeks later, mice were sacrificed for gland function assay and histological analysis.

**Human IFN-γ treatment**. Recombinant human IFN-γ (Peprotech, #300-02) was diluted in 0.1% BSA in PBS. Various concentrations of IFN-γ (20 ng/ml) were used to treat NS-SV-TT-AC cells[13] as for 4 days prior to performing calcium imaging or RNA extraction. IFN-γ was removed 30 min prior to measuring intracellular Ca2+ to lead the cells with Fura2-AM. For RNA extraction, IFN-γ was removed and RNA processing was immediately started.

**Preparation of peripheral blood mononuclear cells (PBMC) for scRNA-Seq**. PBMCs of patients with SjD and non-SjD controls were thawed and stained with Annexin-V (BioLegend, Cat. 640912) and LIVE/DEAD (ThermoFisher, Cat. L23105) in Annexin-V binding buffer (BioLegend, Cat. 422201). Live PBMC were sorted as Annexin-$V^-$ Live/$Dead^-$ on Aria II (BD BioSciences). Cells were collected in RPMI1640 medium (Corning, Cat. 10040CV) with 10% fetal bovine serum. Collected cells were then stained using Type B hashing antibodies (BioLegend), for patient identification post multiplexing (sequences were shown in **Supplementary Table 3**), according to manufacturer's protocol. Collected cells were also quantified using a Bio-Rad TC20 automated cell counter. Stained cells were encapsulated into emulsion droplets using Chromium Controller (10x Genomics) for library prep. scRNA-seq and antibody multiplexing libraries were constructed using Chromium Single Cell 3′ v3.1 Reagent Kit and 3' Feature Barcode Kit (PN-1000268 & 1000262 respectively) according to the manufacturer's protocol. Individual libraries were diluted to 2nM and pooled for sequencing. Pooled batches were sequenced with 100 cycle run kits (28bp Read1, 8bp Index1 and 91bp Read2) on the NovaSeq6000 Sequencing System (Illumina). Amplified cDNA was evaluated with an Agilent BioAnalyzer 2100 using a High Sensitivity DNA Kit (Agilent Technologies). Final libraries were evaluated with an Agilent TapeStation 4200 using High Sensitivity D1000 ScreenTape (Agilent Technologies).

**Single-cell Transcriptome Assembly**. Raw reads from Illumina platform for each batch were converted to Fastq files using Illumina's bcl2fastq package (v2.20). The Fastq files were processed using CellRanger v7.0.0 and aligned to the GRCh38 Genome. Downstream single cell analysis was completed in an R environment (v4.1.1) using Seurat (v4.3).

**Demultiplexing, Normalization and Integration of Samples**. Doublet removal, demultiplexing of samples, and quality control were performed via Seurat HTOdemux function, defining 0.99 quantile as the positive cutoff. Cells found to be positive for more than 1 hashtag were filtered as doublets. Additionally, quality control filters were implanted and were set at ≤ 1000 unique molecular identifiers (UMIs) or ≥ 15%, mitochondrial genes removing any cell that matched either category. A total of 105,152 cells made it through the quality control, with a mean of 6185 cells per sample. Expression levels for the remaining cells were normalized via log normalization and then scaled by 10,000. Each sample was individually integrated using SCT normalization, followed by reciprocal principal-component analysis (PCA) integration, using 4,000 highly variable integration features. Two samples, 1 non-SjD Sicca control and 1 Sjogren's patient, with the highest mean count per cell, were selected as integration anchors; both selected anchors were from the third batch. Batch correction was performed using Harmony (v1.1)[14] with default parameters to regress variation among all 3 batches.

**Cell type clustering and annotation.** $CD4^+$ T cells in PMBC were identified using the integrated assay of all combined samples. We performed PCA, calculating the first 75 components, and generated a uniform manifold approximation map (UMAP) and a t-distributed stochastic neighbor embedding (t-SNE) plot using the first 60 dimensions and 20 nearest neighbors with a minimum distance of 0.3 to visualize the cells. Cell clusters were calculated with the same settings using Louvain clustering, with a resolution of 1.4, resulting in 37 clusters. Clusters were annotated based on cluster markers calculated comparing each cluster to all other clusters using MAST, as implemented in Seurat. 46,376 cells were identified as $CD4^+$ T cells (9 clusters) and were subset for further analysis. We re-scaled, performed variable feature selection, integrated and ran PCA analyses on $CD4^+$ T cells using the same settings described above. In short, a UMAP and a t-SNE plot were generated using the first 60 dimensions, using 15 nearest neighbors to define the neighborhood with a minimum distance of 0.3. New clusters were then calculated using Louvain algorithm integrated in Seurat with the same settings as both dimensional reduction plots, with a resolution of 1.0. A total of 18 clusters were identified and confirmed to have distinct localization in both dimensional reduction plots. Cluster cell type annotation was completed using SingleR, using the Database Immune Cell Expression Data as a reference[15]. Additionally, we calculated cluster markers for all clusters as described above to validate the annotations. One cluster initially labeled as Th1 by SingleR was re-annotated as Tr1 upon discovering canonical markers[16]. Adjacent clusters with matching reference annotations were merged. Two separate memory populations were identified and retained as distinct groups due to their non-adjacent distribution. A total of 12 $CD4^+$ subsets were identified, these cell types were also confirmed using conserved markers for each identified cell type. Canonical markers that were used for confirmation were Naïve (*AK5, LEF1, SATB1, CD55*), Memory (*CD44, IL32, TMSB10*), Th1(*IFNG, CCL4, CXCR5, STAT1*), Th2(*GATA3, DEC2, C-MAF*), Th17(*RORC, IL17A*), Tfh (*TOX, ICOS, CD40L*), CTL (*GZMH, GZMA, GNLY, NKG7*), Treg (*FOXP3, TIGIT, CTLA4, IKZF2*), Tr1(*CTLA4, TIGIT, IFNG, FOXP3(-)*).

**Non-negative matrix factorization (NMF).** Gene expression profiles of different subsets of Sjogren's $CD4^+$ T cells were identified by consensus NMF based on the cNMF implementation of scikit-learn v.20.0[17]. Genes that were not expressed in at least 20 cells across all samples were filtered out, and cells with less than 500 genes were removed). No cells were excluded via this gene number filtering. The top 4000 over-dispersed genes were calculated and used to perform NMF analysis. To determine the most accurate number of components(k), k over a range of 10 to 40 was calculated with each value having 200 iterations. The silhouette score and reconstruction error were calculated as implemented in cNMF, with k=34 identified as the most stable solution. The final consensus solution was determined using a density threshold of 0.1 to exclude outlier solutions. A total of 20 usages were identified that had mean scoring for greater than 0.1 for any cell type. Cell type-related gene expression profiles were evaluated using the mean scores of cells corresponding to their disease states. Marker genes for each GEP were calculated using multiple least squares regression of normalized Z-scored gene expression against the consensus GEP usage matrix as implemented by cNMF. Functional enrichment tests for each GEP were performed using the top 100 marker genes per GEP using the EnrichR platform (v3.2)[18]. cNMF analysis was performed in Python (v.3.7.0) using scanpy (v.1.6.0), pandas (v.1.1.3), numpy (v.1.19.2), matplotlib (v.3.3.2). Visualization was performed in R using the ggplot2 package after conversion into a Seurat object.

**Differential gene expression analysis of scRNA-seq data.** To identify differentially expressed genes (DEG) between Sjogren's patients and non-SjD Sicca controls, we divided each identified cell type into their corresponding disease states. All cell types were confirmed to have representation for all batches and samples. We utilized the MAST algorithm (v1.2.1)[19] implemented by Seurat. Genes were considered significant when found in at least 10% of cells, with an absolute $\log_2$ fold change (LFC) ≥ 0.1, and a Bonferroni-adjusted p-value ≤ 0.1.

**Statistical analyses.** Experimental data analyses were performed using Graph Pad Prism software (v8.0.2). Differences between experimental and control groups were analyzed using Student's *t*-test, and multiple *t*-tests. Bioinformatics analysis was performed in an R statistical environment (v4.1.1). Differences between SjD patients and non-SjD groups as well as *Stim1/2*$^{Foxp3}$ and control mice were analyzed using a

Wilcoxon signed-rank test. Results were considered significant when the p-value was $<0.05$ on two-tailed testing. All results are shown as means ± SD.

**Supplementary Table 1. Primers used for qRT-PCR**

| Gene name | Forward primer | Reverse primer |
|---|---|---|
| *cd4* | CAAGCGCCTAAGAGAGATGG | CACCTGTGCAAGAAGCAGAG |
| *cd8* | ACGGGCATTGCTTCTTCTT | ACAGGGACGAAGCTGACTGT |
| *cd19* | CTGGGACTATCCATCCACCA | TGTCTCCGAGGAAACCTGAC |
| *Gata3* | AGGATGTCCCTGCTCTCCTT | GCCTGCGGACTCTACCATAA |
| *Rorc* | TGCAGGAGTAGGCCACATTACA | GACAGGGAGCCAAGTTCTCA |
| *Il1b* | GGTCAAAGGTTTGGAAGCAG | TGTGAAATGCCACCTTTTGA |
| *Il2* | TGAGCAGGATGGAGAATTACAGG | GTCCAAGTTCATCTTCTAGGCAC |
| *Il4* | GTGTCCTTCTCATGGTGGCT | CAGACATCTTTGCTGCCTCC |
| *Il5* | CCCACGGACAGTTTGATTCT | GCAATGAGACGATGAGGCTT |
| *Il9* | CATCAGTGTCTCTCCGTCCCAACTGATG | GATTTCTGTGTGGCATTGGTCAG |
| *Il10* | GGGTTGCCAAGCCTTATCGGAAAT | CCTTGATTTCTGGGCCATGCTTCT |
| *Il13* | CACACTCCATACCATGCTGC | TGTGTCTCTCCCTCTGACCC |
| *Il17* | CTCCAGAAGGCCCTCAGACTAC | AGCTTTCCCTCCGCATTGACACAG |
| *Il21* | GCCAAACTCAAGCCATCA | AGGAAAGAAACAGAAGCAC |
| *Il35* | GGGATGCCAGAGCACCTG | TCCTAGCCTTTGTGGCTGAG |
| *Tgfb* | TCTCCACTGAGGACACATTGA | ATTCGACATGATCCAGGGAC |
| *Tnfa* | AGGGTCTGGGCCATAGAACT | CCACCACGCTCTTCTGTCTAC |
| *Ifng* | TGAGCTCATTGAATGCTTGG | ACAGCAAGGCGAAAAAGGAT |
| *Stim1* | ATTCGGCAAAACTCTGCTTC | GGCCAGAGTCTCAGCCATAG |
| *Gbp1* | ACATGCCCACAGAAACCCTCCA | AGGCATCTCGTTTGGCTTCCAG |
| *Stat1* | GCCTCTCATTGTCACCGAAGAAC | TGGCTGACGTTGGAGATCACCA |
| *Cxcl9* | CCTAGTGATAAGGAATGCACGATG | CTAGGCAGGTTTGATCTCCGTTC |
| *P2RX7* | TAACTGGAACAGAGGGCAAG | CATTGGGTAGCTCAGCAGTT |
| *CHRM3* | TGGACGATGGAGGCAGTTTT | TGAAGGCAAGCAAGATCGCA |
| *GAPDH* | GACAGTCAGCCGCATCTTC | GCGCCCAATACGACCAAAT |

**Supplementary Table 2. Reagents used for flow cytometry**

| Mouse Antigen | Clone | Conjugated FC | Source | Dilution |
|---|---|---|---|---|
| CD45 | 30-F11 | PerCP-Cy5.5 | BioLegend | 1:400 |
| CD3 | 17A2 | PE-Cy7 | BioLegend | 1:400 |
| CD4 | GK1.5 | APC/Cy7 | BioLegend | 1:500 |
| CD8a | RM4-5 | PerCP-Cy5.5 | BioLegend | 1:400 |
| B220 | RA3-6B2 | BV510 | BioLegend | 1:400 |
| CD11b | M1/70 | PE-Cy7 | BioLegend | 1:500 |
| Foxp3 | FJK-16s | PE | eBioscience | 1:100 |
| Foxp3 | FJK-16s | FITC | eBioscience | 1:100 |
| CD25 | PC61 | APC | BioLegend | 1:500 |
| CXCR3 | CXCR3-173 | PE | BioLegend | 1:400 |
| IFN-γ | XMG1.2 | PE | BioLegend | 1:200 |
| TNF-α | MP6-XT22 | APC | eBioscience | 1:200 |
| CD95(Fas) | Jo2 | BV421 | BD Bioscience | 1:200 |
| CD138 | 281-2 | PE | Biolegend | 1:1000 |
| GL-7 | GL7 | FITC | Biolegend | 1:1000 |
| CD38 | T10 | PE-Cy7 | Biolegend | 1:500 |
| GATA3 | TWAJ | eFluor® 660 | eBioscience | 1:100 |
| T-bet | eBio4B10 | PE | eBioscience | 1:100 |
| Annexin-V | N/A | APC | BioLegend | 1:100 |
| Live/Dead | N/A | Blue | Thermo Fisher | 1:500 |

### Supplementary Table 3. Patient information and criteria.

| Subject id | Age | Gender | Anti-SS-A | ACR/ EULAR Score* | Focus Score | Baseline UWS flow rate | Baseline Max Ocular Staining score >=3 | Diagnosis | Group | Hash-TAG Seq |
|---|---|---|---|---|---|---|---|---|---|---|
| 773251 | 66 | Male | Negative | 1 | N/A | UWS < 0.5 | No | Non-specific chronic inflammation | control | TGATGGCCTATTGGG |
| 770586 | 59 | Female | Negative | 1 | 0.3 | UWS < 0.5 | No | Focal lymphocytic sialendenitis | control | TTCCGCCTCTCTTTG |
| 770575 | 63 | Female | Negative | 5 | 1.9 | UWS < 0.5 | Yes | Focal lymphocytic sialendenitis | pSS | TGTCTTTCCTGCCAG |
| 770272 | 71 | Female | Positive | 9 | 12 | UWS < 0.5 | Yes | Focal lymphocytic sialendenitis | pSS | AGTAAGTTCAGCGTA |
| 770400 | 28 | Male | Positive | 9 | 6.6 | UWS < 0.5 | Yes | Focal lymphocytic sialendenitis | pSS | AAGTATCGTTTCGCA |
| 772193 | 40 | Female | Negative | 0 | 0.3 | UWS >= 0.5 | No | Focal lymphocytic sialendenitis | control | AAGTATCGTTTCGCA |
| 770784 | 60 | Female | Negative | 1 | 0.8 | UWS < 0.5 | No | Focal/sclerosing lymphocytic sia | control | GGTTGCCAGATGTCA |
| 772246 | 57 | Female | Negative | 1 | N/A | UWS < 0.5 | No | Sclerosing chronic sialadenitis | control | TGTCTTTCCTGCCAG |
| 770611 | 67 | Female | Positive | 7 | 4.6 | UWS >= 0.5 | Yes | Focal/sclerosing lymphocytic sia | pSS | CTCCTCTGCAATTAC |
| 772215 | 28 | Female | Positive | 8 | 2.2 | UWS < 0.5 | Yes | Focal lymphocytic sialendenitis | pSS | CAGTAGTCACGGTCA |
| 773261 | 79 | Female | Positive | 9 | 2.5 | UWS < 0.5 | Yes | Focal lymphocytic sialendenitis | pSS | ATTGACCCGCGTTAG |
| 772456 | 54 | Female | Negative | 0 | N/A | UWS >= 0.5 | Yes | Sclerosing chronic sialadenitis | control | AAGTATCGTTTCGCA |
| 770287 | 64 | Female | Negative | 1 | 0.5 | UWS >= 0.5 | Yes | Focal lymphocytic sialendenitis | control | GGTTGCCAGATGTCA |
| 772236 | 45 | Female | Negative | 0 | 0.3 | UWS >= 0.5 | Yes | Focal lymphocytic sialendenitis | control | TGTCTTTCCTGCCAG |
| 772421 | 54 | Female | Positive | 6 | N/A | UWS < 0.5 | Yes | Sclerosing chronic sialadenitis | pSS | CTCCTCTGCAATTAC |
| 770399 | 48 | Female | Positive | 8 | 3.7 | UWS < 0.5 | Yes | Focal lymphocytic sialendenitis | pSS | CAGTAGTCACGGTCA |
| 772530 | 44 | Female | Positive | 9 | 1.4 | UWS < 0.5 | Yes | Focal lymphocytic sialendenitis | pSS | ATTGACCCGCGTTAG |

Abbreviations: pSS, primary Sjogren's syndrome; UWS, unstimulated whole saliva flow.

**Supplementary Table 4. List of genes regulating $Ca^{2+}$ homeostasis**

| | | | | | | |
|---|---|---|---|---|---|---|
| *CRACR2A* | *JPH3* | *ORAI2* | *STIM2* | *ATP2A3* | *P2RX1* | *MCU* |
| *CRACR2B* | *JPH4* | *ORAI3* | *TMEM110* | *ATP2B1* | *P2RX2* | *MCUB* |
| *ITPR1* | *KCNA3* | *RYR1* | *TRPM4* | *ATP2B2* | *P2RX3* | *SMDT1* |
| *ITPR2* | *KCNN4* | *RYR2* | *PGAM5* | *ATP2B3* | *P2RX4* | *MICU1* |
| *ITPR3* | *MBP* | *RYR3* | *LRRC8C* | *ATP2B4* | *P2RX5* | *MICU2* |
| *JPH1* | *NAPA* | *SARAF* | *ATP2A1* | *ATP2C1* | *P2RX6* | *MCUR1* |
| *JPH2* | *ORAI1* | *STIM1* | *ATP2A2* | *ATP2C2* | *P2RX7* | *SLC8B1* |

Supplementary Figure 1

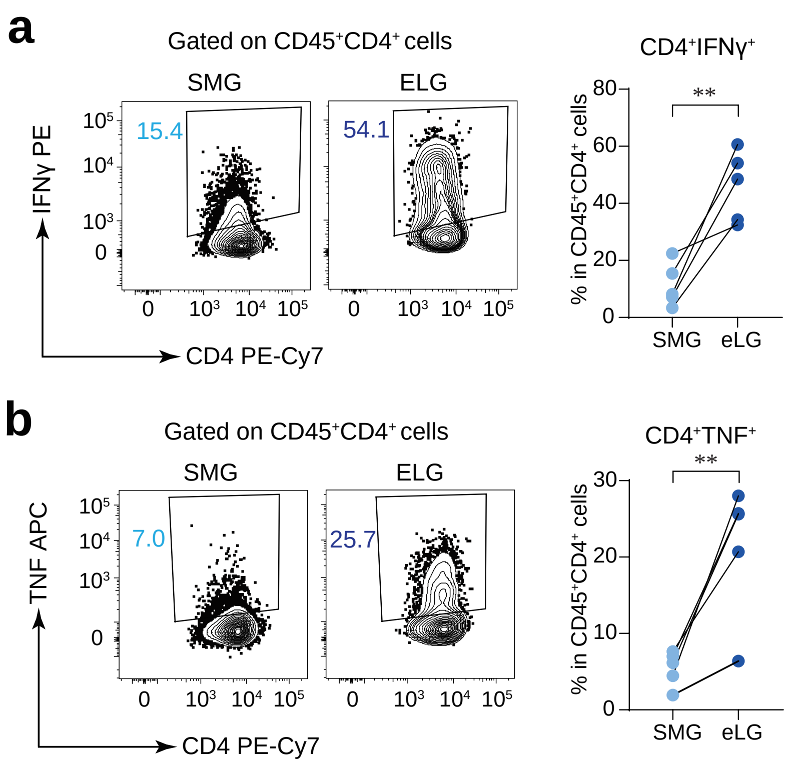


**Supplementary Figure 1. Lacrimal gland derived T cells produce IFNγ and TNF in *Stim1/2$^{Foxp3}$* mice.** **a, b)** Representative flow cytometry plots and bar graphs showing the frequencies of $CD4^{+}IFN\gamma^{+}$ **(a)** and $CD4^{+}TNF^{+}$ **(b)** T cells in SMG and ELG from male Cre-negative littermate control and *Stim1/2$^{Foxp3}$* mice after 4h restimulation with PMA + Ionomycin. Data are from 5 mice per cohort with 1 technical replicate per mouse. Statistical analyses were conducted using two-tailed, paired Student's t test, and shown as means ± SD. *$P$ < 0.05, **$P$ < 0.01 and ***$P$ < 0.001.

Supplementary Figure 2

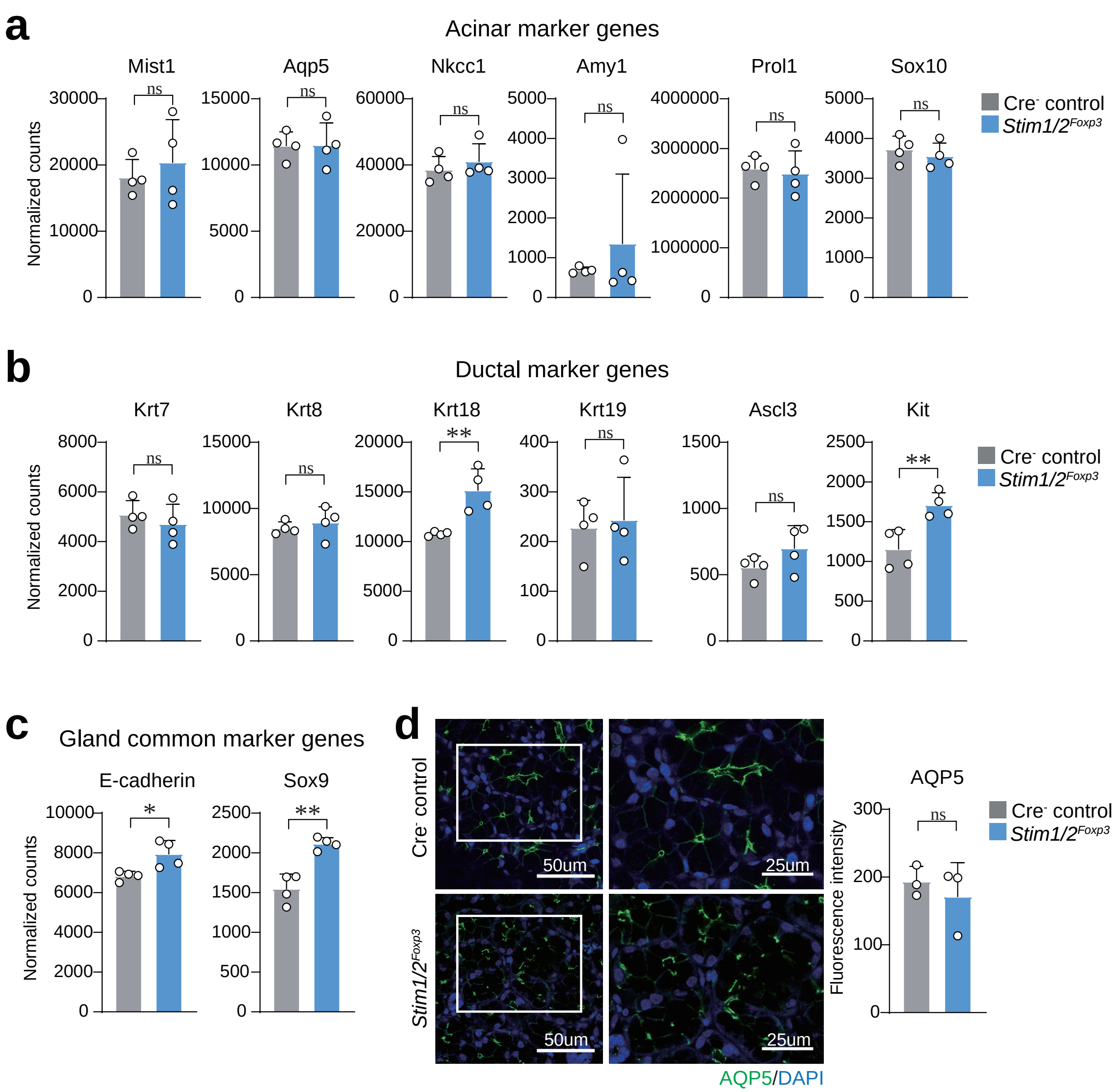


**Supplementary Figure 2. Comparable gland marker gene expression and AQP5 level in salivary gland of *Stim1/2$^{Foxp3}$* mice.**

**a–c)** Normalized count of acinar marker genes **(a)**, ductal marker genes **(b)** and gland common marker genes **(c)** in transcriptomes analysis of SMG isolated from male Cre-negative littermate control and *Stim1/2$^{Foxp3}$* mice by RNA-sequencing. Data are from 4 mice per cohort with 1 technical replicate per mouse. **d)** Representative immunofluorescence images and bar graphs showing the quantified intensity of AQP5 staining in SMG from male Cre-negative littermate control and *Stim1/2$^{Foxp3}$* mice. Data are from 3 mice per cohort with 1-6 images per mouse. Statistical analyses were conducted using two-tailed, unpaired Student's t test, and shown as means ± SD. **P* < 0.05, ***P* < 0.01 and ****P* < 0.001.

Supplementary Figure 3

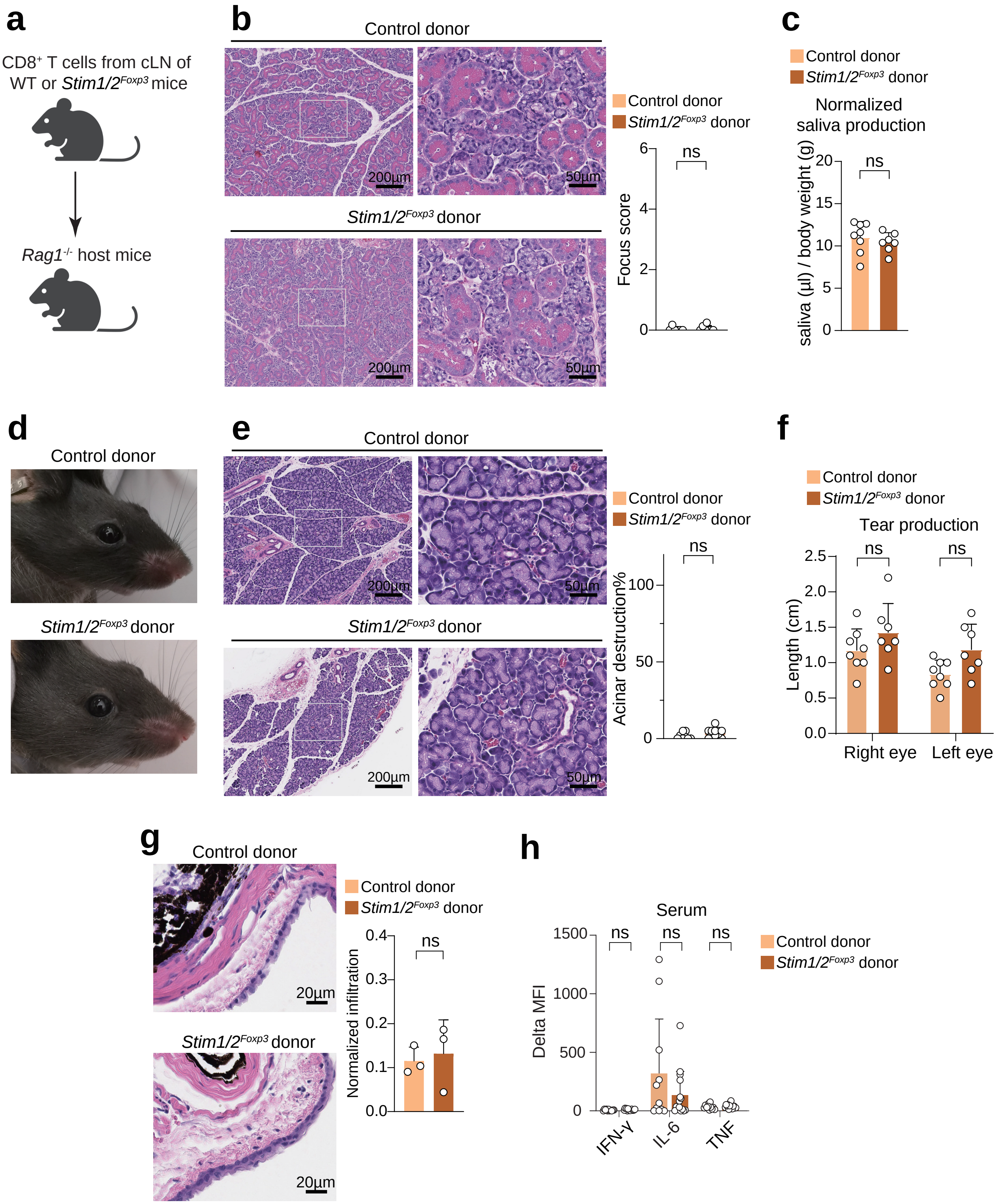

**Supplementary Figure 3. CD8$^{+}$ T cells transfer does not induce gland inflammation and phenotypes of SjD in *Rag1$^{-/-}$* mice.**

**a)** Design of experiments shown in panels **b–h**. CD8$^{+}$ T cells were isolated from cLN of male *Stim1/2$^{Foxp3}$* and Cre-negative littermate control mice, and injected i.v. into male *Rag1$^{-/-}$* mice. **b)** Representative H&E staining and modified focus scores of SMG from *Rag1$^{-/-}$* host mice. Data are from 6-7 mice per cohort with 1 gland (right) per mouse. **c)** Saliva production over a period of 20 min after pilocarpine stimulation of *Rag1$^{-/-}$* host mice. Data are from 7-8 mice per cohort with 1 measurement per mouse. **d, e)** Representative ocular images, H&E staining and quantified exorbital lacrimal gland (ELG) destruction in *Rag1$^{-/-}$* host mice. Data are from 8-11 mice per cohort with 1 eye or gland (right) per mouse. **f)** Analysis of tear production by phenol red thread test. Data are from 7-8 mice per cohort with 1 measurement per eye. **g)** Representative H&E staining of conjunctiva and quantified conjunctivitis from *Rag1$^{-/-}$* host mice. Data are from 3 mice per cohort with 2 eyes per mouse. **h)** Analysis of IFN-γ, TNF, and IL-6 concentration in the serum of *Rag1$^{-/-}$* host mice by CBA, Data are from 11 to 15 mice per cohort with 1 technical replicate per mouse. Statistical analyses were conducted using two-tailed, unpaired Student's *t*-test, and shown as means ± SD. *$P < 0.05$, **$P < 0.01$ and ***$P < 0.001$.

Supplementary Figure 4

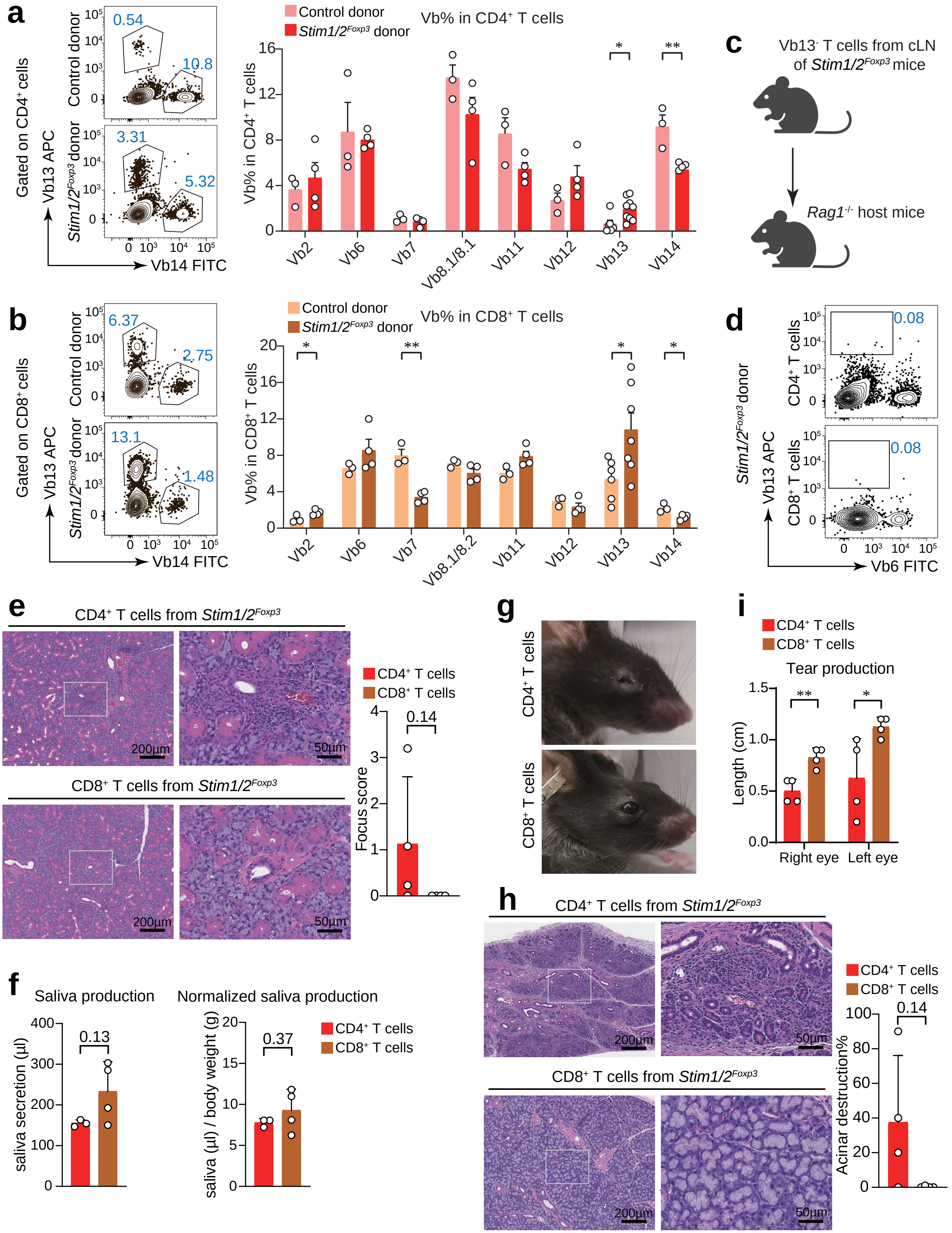

**Supplementary Figure 4. Vβ13$^{-}$CD4$^{+}$ T cells are sufficient to induce phenotypes of SjD in *Rag1$^{-/-}$* mice.**

**a, b)** Representative flow cytometry plots and bar graphs showing the T cell V beta chain usage in cLN of *Rag1$^{-/-}$* host mice received CD4$^{+}$CD25$^{-}$ T cells and CD8$^{+}$ T cells from Cre-negative littermate control and *Stim1/2$^{Foxp3}$* mice. Data are from 3-9 mice per cohort from 2 independent experiments with 1 technical replicate per mouse. **c)** Design of experiments shown in panels **d–i**. Vβ13$^{-}$CD4$^{+}$CD25$^{-}$ and Vβ13$^{-}$CD8$^{+}$ T cells were isolated from cLN of male *Stim1/2$^{Foxp3}$* mice, and injected i.v. into male *Rag1$^{-/-}$* mice respectively. **d)** Representative flow cytometry plots showing the frequencies of Vβ13$^{+}$CD4$^{+}$ and Vβ13$^{+}$CD8$^{+}$ T cells in cLN of *Rag1$^{-/-}$* host mice received CD4$^{+}$CD25$^{-}$ T cells and CD8$^{+}$ T cells from *Stim1/2$^{Foxp3}$* mice. Data are from 4 mice per cohort with 1 technical replicate per mouse. **e)** Representative H&E staining and modified focus scores of SMG from *Rag1$^{-/-}$* host mice. Data are from 4 mice per cohort with 1 gland (right) per mouse. **f)** Saliva production over a period of 20 min after pilocarpine stimulation of *Rag1$^{-/-}$* host mice. Data are from 3-4 mice per cohort with 1 measurement per mouse. **g, h)** Representative ocular images, H&E staining and quantified exorbital lacrimal gland (ELG) destruction in *Rag1$^{-/-}$* host mice. Data are from 4 mice per cohort with 1 eye or gland (right) per mouse. **i)** Analysis of tear production by phenol red thread test. Data are from 4 mice per cohort with 1 measurement per eye. Statistical analyses were conducted using two-tailed, unpaired Student's *t*-test, and shown as means ± SD. *$P < 0.05$, **$P < 0.01$ and ***$P < 0.001$.

Supplementary Figure 5

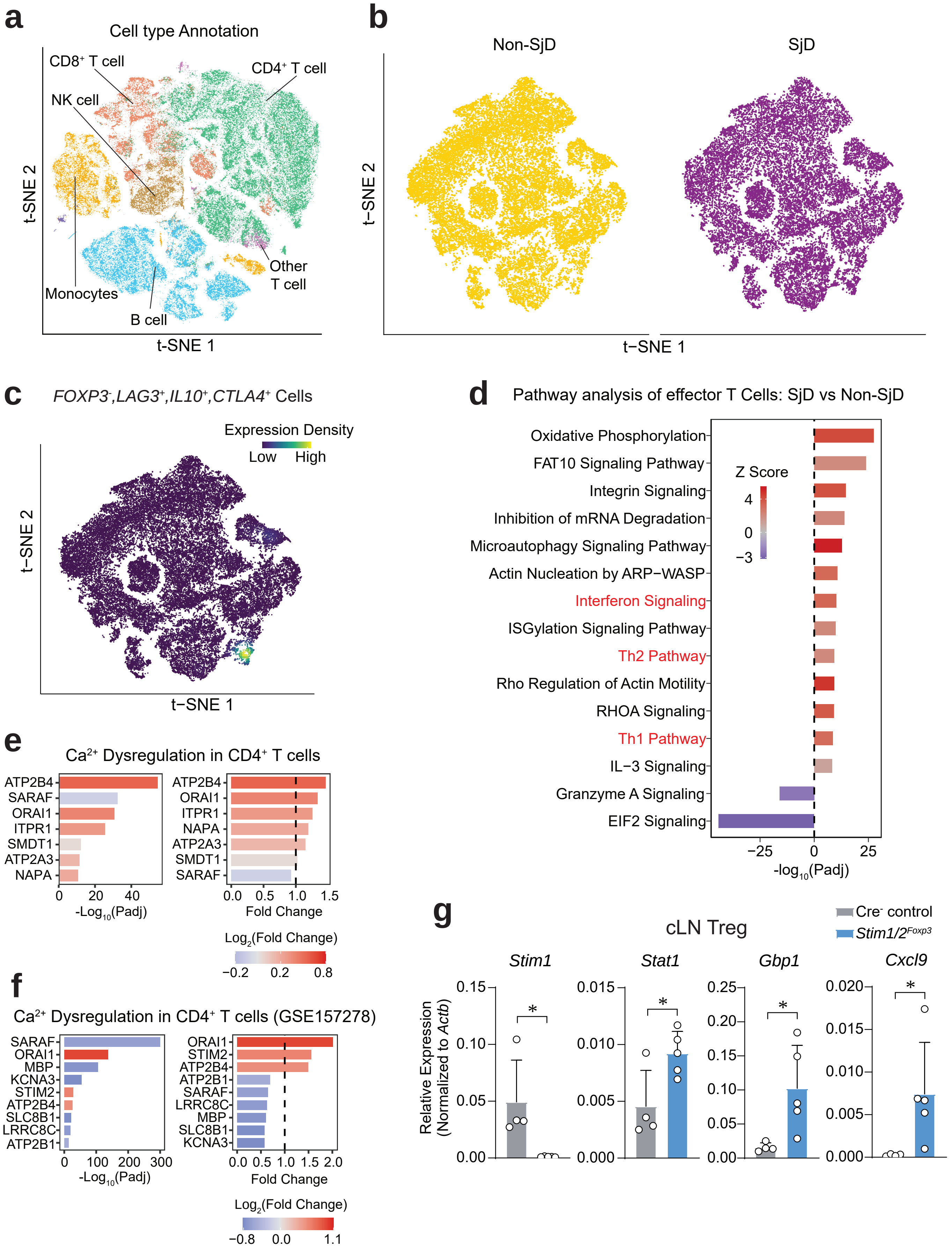

**Supplementary Figure 5. Patients with SjD have increased Treg cell frequency but enhanced Th1 and Th2 differentiation based on single-cell RNA-seq of PBMCs.**

**a)** Aggregate t-SNE reduction of 105,000 cells from human PBMCs of 9 patients with SjD and 8 symptomatic non-SjD controls. $CD4^+$ T cells were identified by marker gene expression. **b)** t-SNE reduction of 46,376 $CD4^+$ T cells from human PBMCs of 9 patients with SjD and 8 symptomatic non-SjD controls. **c)** t-SNE representation of $CD4^+$ T cells expressing Tr1 marker genes *(CTLA4, LAG3, IL10, FOXP3(-))* from 9 patients with SjD and 8 symptomatic non-SjD controls. **d)** Enrichment pathway analysis for effector $CD4^+$ T cells using DEGs based on IPA database. Unbiased top 15 dysregulated pathways were ranked by *P* value (-log10). **e,f)** Differentially expressed genes (DEG) that regulate $Ca^{2+}$ homeostasis in all $CD4^+$ T cells from 9 patients with SjD and 8 symptomatic non-SjD controls **(e),** and 5 patients with SjD patients and 5 controls using data from GSE157278 **(f)**. **g)** mRNA expression of interferon stimulated gene in Treg cells from cLN of male *Foxp3*-GFP reporter control and $Stim1/2^{Foxp3}$ mice measured by RT-qPCR. RT-qPCR data were normalized to *Actb* housing control expression. Data are from 4-5 mice per cohort with 3 technical replicates per mouse. Statistical analysis in panel d-f was completed using Wilcoxon rank-sum test, and significance was adjusted using the Benjamin Holmberg method. And panel g was analyzed by two-tailed, unpaired Student's *t*-test. Results shown as means ± SD. $*P < 0.05$, $**P < 0.01$ and $***P < 0.001$.
